# Spatial segregation and multi-scale space-use adjustments to socio-ecological variables in western gorillas

**DOI:** 10.64898/2026.10.19.759586

**Authors:** Benjamin Robira, Simon Benhamou, Thomas Breuer, Terence Fuh, Erlich Obeki Bayanga, Silvia Miglietta, Andréa Sotto-Mayor, Giulia Bardino, Chloé Cipolletta, Angelique Todd, Shelly Masi

**Affiliations:** CEFE, Université de Montpellier & CNRS, Montpellier, France; Eco-anthropologie (EA), Muséum National d’Histoire Naturelle, CNRS, Université Paris Cité, Musée de l’Homme, Paris, France; Associated to Cogitamus Lab; Wildlife Conservation Society, Global Conservation Program, New-York, USA; World Wide Fund for Nature Germany, Berlin, Germany; Dzanga-Sangha Protected Areas, WWF, Bangui, Central African Republic; Mondika Research Center; Nouabalé-Ndoki National Park, Wildlife Conservation Society; Congo Program, Republic of Congo; Department of Biology and Biotechnology “Charles Darwin” (BBCD), University of Rome “La Sapienza,” Rome, Italy

**Author notes:** Institute for Biodiversity and Ecosystem Dynamics, University of Amsterdam, Amsterdam, Netherlands (Benjamin Robira, Shelly Masi). Correspondence: Shelly Masi < >; **<u>Benjamin Robira</u>** < >.

**Keywords:** Daily movement, *Gorilla gorilla*, Home range, Indirect competition, Overlap, Site fidelity, Spatial segregation

## Abstract

**Background:** Identifying the mechanisms that shape space-use dynamics and the scale at which they operate can provide insights into the socio-ecology of numerous animal species. This is crucial for addressing both applied (e.g., conservation) and theoretical questions (e.g., the coexistence or segregation of individuals). Yet, longitudinal views of the space-use dynamics are generally rare. Taking advantage of a long-term movement dataset (up to 8 years) on five habituated groups of wild western gorillas (*Gorilla gorilla*) in Central Africa, we investigated the dynamics of their space use in relation to the socio-ecological variables from everyday movement to home range shifts.

**Methods:** We looked at which temporal scale their ranging movements are stationary to highlight the dynamics of home range settling and shifts. We also studied how daily space use is shaped by (i) foraging and intra-group constraints (i.e., diet, group size, presence of offspring), and by (ii) inter-group competition (direct or indirect), quantifying static and dynamic interactions in neighbouring group-dyads.

**Results:** Gorilla groups shifted the centre of their home range 0.17 times per year by a distance larger than the mean home range radius. Movements within the home range were highly dependent on the group composition and the ecological features (with larger daily-travelled distances in the high-fruit season compared to the low-fruit season). The presence of dependent infants affected the daily-travel distance: the larger their number, the smaller the daily-travel distance. Furthermore, the frequently visited sites of the home range were also used the longest, but their locations differed seasonally, suggesting space use flexibility. Although there was little overlap between the home ranges of neighbouring groups, these groups did not seem to avoid each other more than expected at random. They could even venture into, and slept in, the central part of the home range of a neighbouring group.

**Conclusion:** Our study suggests that spatial segregation should be the simple consequence of avoidance of depleted areas rather than of territorial behaviour. It shows how socio-ecological constraints may shape the space-use patterns at different scales in a non-territorial species, and questions the role of cognition in space-use dynamics.

## BACKGROUND

The locomotor abilities, physiology (e.g., body condition, Goossens et al. 2020), and cognitive level of an animal (in particular, spatial memory; Fagan et al. 2013), as well as the ecological challenges it faces (e.g., food distribution, Bartumeus et al. 2005), all contribute to shaping its movements. For group-living animals, movement is also further affected by social factors, such as within-group cohesion and competition, and inter-group interactions (Shaw 2020). These different facets of animal movement match with the plethora of space-use strategies, from nomadism to sedentarity (Teitelbaum and Mueller 2019).

In most mammal species, individuals restrict their movements within limited areas called home ranges, which are much smaller than expected based on their movement abilities alone (Burt 1943; Börger, Dalziel, and Fryxell 2008; Powell and Mitchell 2012). Large differences in space-use patterns and dynamics at the home range scale may exist even in closely related species (e.g., for primates: Herbinger, Boesch, and Rothe 2001; Watts and Mitani 2001; Langergraber et al. 2017; Samuni, Wegdell, and Surbeck 2020), dramatically limiting possible extrapolation, even to sister species. Highlighting a species’ space-use dynamics is yet essential to understand its socioecology (Börger, Dalziel, and Fryxell 2008) and population dynamics (Morales et al. 2010; Barraquand and Murrell 2012; Riotte-Lambert et al. 2017). This thus provides key knowledge for assisting conservation plans, particularly for endangered species.

Animals that display site fidelity through home range behaviour can become highly familiar with their local environment (Piper 2011). As such, site fidelity is often associated with a higher foraging efficiency (Riotte-Lambert, Benhamou, and Chamaillé-Jammes 2015) and a reduced risk of predation (Gehr et al. 2020). However, it may be disrupted by seasonal changes in food availability (e.g., Cornélis et al. 2011, Avgar, Street, and Fryxell 2014) or by social and reproductive events (e.g., dispersal at sexual maturity, Bowler and Benton 2005).

Site fidelity is more beneficial when an animal or a group can use the area exclusively (i.e., without intra-specific competition). Exclusive use of space may occur through territorial behaviour, whereby animals actively defend their home ranges or parts of them to gain exclusive access to food and/or mates (Burt 1943; Grant, Chapman, and Richardson 1992). Territoriality is generally expressed through aggressive behaviours during group encounters, but also through animals’ movements (e.g., through border patrolling, Langergraber et al. 2017). In primates, the daily path length of territorial species has been shown to be larger than one would expect based on their home range size, all else being equal (Mitani and Rodman 1979). Conversely, exclusive space use may also occur passively as a by-product of spatial memory, due to foraging optimisation by memorising and avoiding areas depleted by neighbouring conspecifics (Riotte-Lambert, Benhamou, and Chamaillé-Jammes 2015; Aarts et al. 2021; Ranc, Cagnacci, and Moorcroft 2021),

The social and ecological factors affecting movement and site exclusion mechanisms are central to understand spatial cognition in numerous species. On the one hand, patterns of food availability should imply weak to intensive selection towards higher cognition, depending on whether the distribution is homogeneous or heterogeneous in space and time (*Ecological Intelligence Hypothesis*; Milton 1981; Rosati 2017). On the other hand, the positive correlation between home range overlap and brain size (Grueter 2015) has been considered as supporting the hypothesis that intergroup interactions should shape selection towards higher cognition (*Napoleonic Intelligence Hypothesis*; Ashton, Kennedy, and Radford 2020). As such, the characterisation of socio-ecological factors affecting the dynamics of space use can also provide helpful insights on the evolution of animal cognition (e.g., see Janmaat and Chancellor 2010).

The western gorilla (*Gorilla gorilla*) is an interesting species to investigate the socio-ecological factors affecting movement patterns. They differ greatly from those of the much more widely studied mountain gorilla (*Gorilla beringei beringei*). First, western gorillas rely on fruit, a highly scattered and ephemeral resource (Janmaat et al. 2016). Their space use behaviour is thus driven by the periodic oscillation of fruit availability: they travel longer distances, over longer durations, and cover larger areas (Remis 1997; Goldsmith 1999; Cipolletta 2004; Masi, Cipolletta, and Robbins 2009; Seiler and Robbins 2020; Robira et al. 2023a) during periods of high fruit abundance, when compared to periods of fruit scarcity. Second, groups of western gorilla include only one dominant male (Forcina et al. 2019; Masi et al. 2021; vs. several ones in mountain gorillas, Robbins, 1995; Seiler et al. 2017), rendering western gorilla particularly sensitive to group encounters due to risk of infanticide and/or the transfer of females (Harcourt and Stewart 2007; Potel et al. 2023). Despite recent investigations, a comprehensive understanding of how these factors together influence the general movement strategy and associated space use patterns of western gorillas, and the timescale over which this occurs, is lacking. Indeed, most studies on western gorillas have focused separately either on the social or the ecological factors of space use. Such studies were furthermore subject to biases due to the monitoring approach, as they either relied on indirect tracking methods (via camera traps, Morrison et al. 2020), or on live tracking through handheld GPS of the human trackers, but on groups not yet fully habituated to human presence (Cipolletta 2003, 2004). The long-term tracking of a handful of groups has recently enabled a more thorough characterization of western gorilla space-use patterns (Judson et al. 2023; Scarry et al. 2023). However, these studies remained limited in their space use quantification and number of study groups.

Using a unique long-term dataset on movement data of five habituated groups of free-ranging western gorillas (including two neighbouring dyads), here we aimed to understand how the space-use patterns result from individual needs, food distribution, spatial memory as well as from potential interaction with other groups. In particular, we aimed to elucidate the roles played by ecological (e.g., seasonal diet) and social (e.g., group size, interaction with neighbouring groups) factors on their space-use at different scales, from everyday movements to home range shifts. In this framework, we investigated different space-use variables (daily path length, home range size, site fidelity, recursion and intensity distributions, space-use overlap between neighbouring groups) to address the following questions:

- To what extent do home range size (as a proxy for space defensibility), group size (as a proxy for energetic requirements and food competition) and season (as a proxy for environmental complexity) affect daily path length? As found in other animal species, including primates (Caillaud et al. 2014; Ganas and Robbins 2005), we expected that the daily path length would be positively affected by both the home range area and the group size, in addition to the increase in fruit consumption (e.g., Cipolletta, 2002, 2003).
- How much are western gorillas faithful to an area and what factors are associated with site fidelity disruption? Given that these animals strongly rely on memory to forage (Salmi et al. 2020; Robira et al. 2023a, b; Robira et al. 2024), and may thus benefit a lot from familiarity with their local environment, we expected them to shift their home ranges only rarely. Furthermore, because inter-group dynamics in western gorillas can lead to long-term and extremely potential high risk (infanticides and female transfers, Harcourt and Stewart 2007; Potel et al. 2023), we predicted that shifts should be triggered more by social factors (e.g., vicinity of other groups) than by ecological ones (e.g., seasonal changes in the diet, for which it is advantageous to know the area well in order to find food).
- Do neighbouring groups segregate in space, and what mechanisms are likely to drive the level of space sharing? Western gorillas have recently been claimed to be territorial, based on evidence of avoidance provided by indirect monitoring by camera traps (Morrison et al. 2020). To question further this finding, we investigated static (home range overlap, asymmetry of the use intensity of overlapping areas) and dynamic (simultaneous movement) interactions between neighbouring groups based on direct simultaneous tracking, and the relationship between daily-travel distance and home range size. Territoriality should be evidenced by low overlap, simultaneous avoidance and longer daily-travel distances than expected based on the home range size. Should the latter be wrong and space-use asymmetry low, then passive segregation due to memory would be more likely.

## METHODS

### Study sites and gorilla groups

This study was carried out at three field research stations within the Sangha Tri-National Landscape (locally known by its French acronym TNS, Figure 1): Bai Hokou (N 2° 52’, E 16° 28’), Mongambe (N 2° 55’; E 16° 23’), both in the Dzanga-Ndoki National Park in the south-western part of the Central African Republic (CAR), and Mondika (N 4° 39’; E 18° 56’) in the periphery of the Nouabalé-Ndoki National Park in the Republic of Congo (RC). The TNS environment has a seasonal climate with < 100 mm monthly rainfall in December-February and > 150 mm monthly rainfall in September-October. The region is characterised by a tropical rainforest which is a mosaic of primary monodominant *Gilbertiodendron* forest and mixed forest, including secondary *Marantaceae* forest patches, swamps, and natural forest clearings. Data were collected on five human-habituated groups of wild western gorillas: CAR1 (7 individuals) and CAR3 (8 to 9) from Bai Hokou, and CAR2 (9 to 10) from Mongambe, and RC1 (13 to 14) and RC2 (6 to 11) from Mondika. Group compositions and their eventual changes during the study periods are in Table S1 of the Supplementary Material. While CAR1, CAR2, RC1, and RC2 groups were fully habituated to close human presence, CAR3 was at the latest stage of habituation (with the majority of the individuals tolerating human presence up to about 10 m).

**Figure 1.**
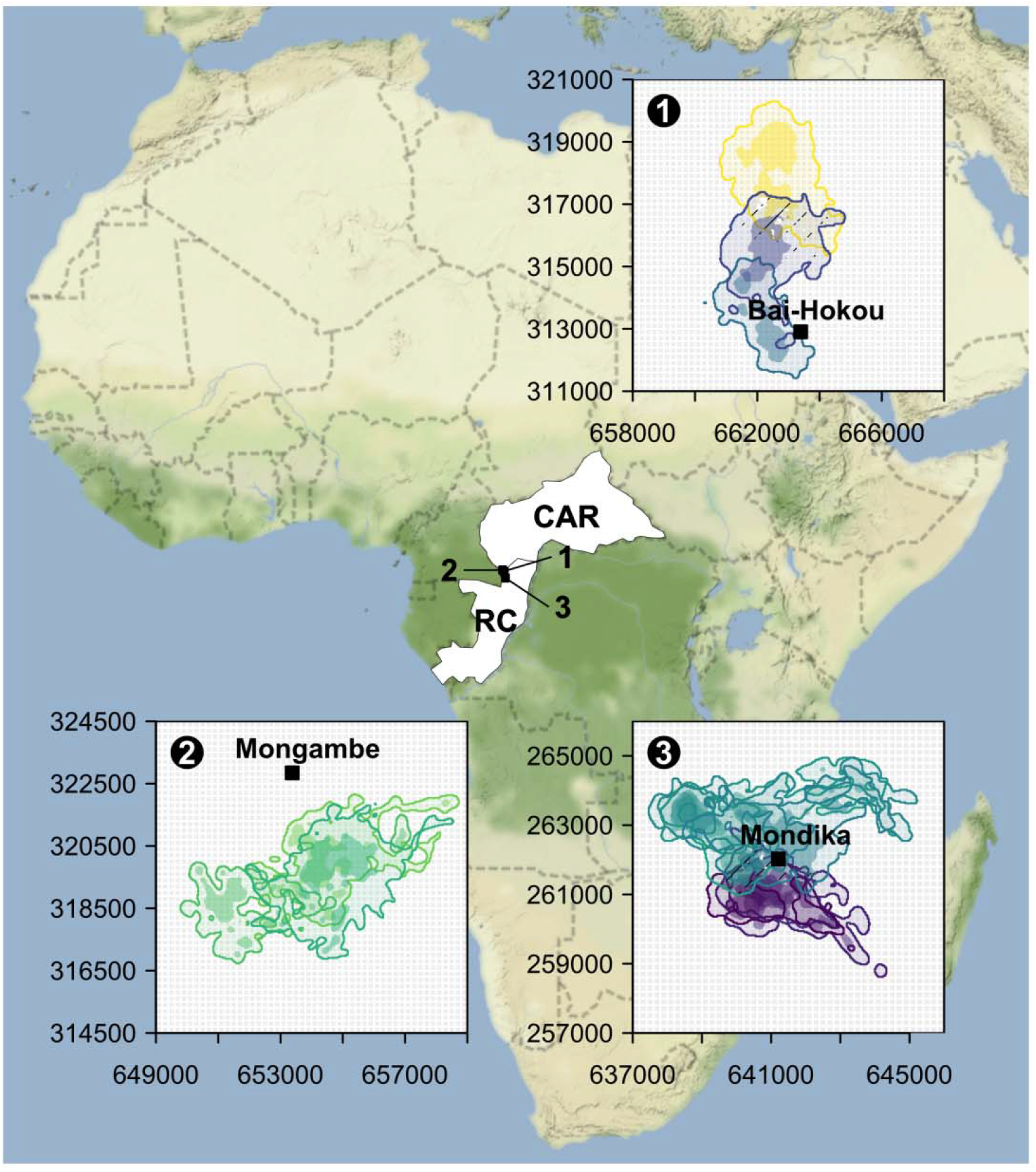
Study sites and study groups’ home ranges | The home range areas and the corresponding core areas (within the 95% and 50% cumulative isopleths of the UDs, respectively) are shown with light and dark colours, respectively. Camp sites are indicated by black squares. They are ca. 30 km away from each other, on each side of the border between the Central African Republic (CAR) and the Republic of the Congo (RC). The spatial projection corresponds to the UTM 33N.

### Data recording

We tracked gorilla groups from sunrise to sunset by following them at a very short distance. During a first tracking period, which involved only CAR1 (first tracked from February 2003 to August 2004, and then from January 2009 to September 2014, totalling 1904 days) and CAR2 (tracked from January 2008 to August 2014, totalling 2194 days), locations were recorded based on a grid-system made of 250 m side-quadrats, with the help of a compass and an extensive network of mapped trails (see Cipolletta 2003, 2004; note that > 1-month gaps occurred because of data loss for CAR2 group because of a fire at the camp). During a second tracking period, from June 2016 to November 2017, CAR1, CAR2, and CAR3, were simultaneously GPS-tracked (using a handheld GPS receiver) for 469, 380 and 337 days, respectively. A new GPS location was recorded every 18 meters (the minimum recording distance allowed in the track-log setting). RC1 and RC2 were also simultaneously GPS-tracked from October 2012 to January 2015, for 596 and 523 days, respectively, but with relocations recorded every 20 min. While tracking gorillas, we also daily collected information about the group composition (presence of newborn or immigrating individuals, deaths or emigrations of individuals) and possible interactions with other social units (other groups or solitary males). The occurrences of such interactions were detected based on direct (visual or auditory) or indirect observations of other gorillas (traces along the gorilla trails or at the nest sites), or on alert reactions from the silverback (e.g., chest beating, hooting, slap on ground/tree) or other group members (e.g., hand clapping, screaming, running away etc.). Age-sex classes were defined following Breuer et al. (2009).

### Analyses

#### DATA processing and analyses were implemented in the *R* environment (v.4.1.2, R Core Team 2020)

##### Locational stationarity

Home-range behaviour involves the absence of significant changes in mean and/or variance in location across time (i.e., locational stationarity) over a period of at least several weeks, due to repeated visits to the same restricted set of places. If the location time series over a large temporal extent (e.g., several years) is stationary, a long-term home range can be defined. Additionally, irrespective of whether the series is stationary or not in the long-term, it may be piecewise stationary, i.e., made of phases that are stationary over shorter periods (e.g. a few weeks), corresponding to temporary (e.g., seasonal) home ranges (Benhamou 2014) that are thus highlighted on a biological basis rather than on an arbitrary time basis (e.g., monthly, as often done). We looked at possible global stationarity and at piecewise stationarity with phases > 30 days using the “segmentation” function of the *segclust2d* package (Patin et al. 2020) on the bivariate time series of easting (*x*) and northing (y) coordinates. For the *grid data set*, we used the coordinates of the quadrat centre (see Figure 2 for example) with a timestamp defined only on a daily basis. For the *GPS data set*, we kept a GPS location every 2 hours between 8:00 to 16:00 (gorillas are strictly diurnal) as keeping more than five locations a day may result in computer memory issues. The output of the segmentation of the location time series obtained from the *GPS data set* is shown in Supplementary Material 3. To quantify the overall tendency of a group to change its home range (change in location and/or in size), we calculated the Simpson (1949) index, which ranges between 0 and 1, on the relative time lengths of each stationary phase 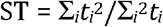, where *t*_i_ is the duration of the *i*^th^ stationary phase. A ST value larger than 0.25 is assumed to indicate the occurrence of a major stationary phase. However, with our data, this index has to be interpreted with caution for two reasons. (1) It may be biased by the occurrences of time gaps (i.e., long periods without tracking; this happened only for the *grid data set* because the group was lost; Figure 2; for the *GPS data set*, time gaps were always lower than 22 days). (2) The ST index is also affected by the arbitrary beginning and end of the tracking periods (the first and last stationary phases should have lasted more or less than their apparent durations).

**Figure 2.**
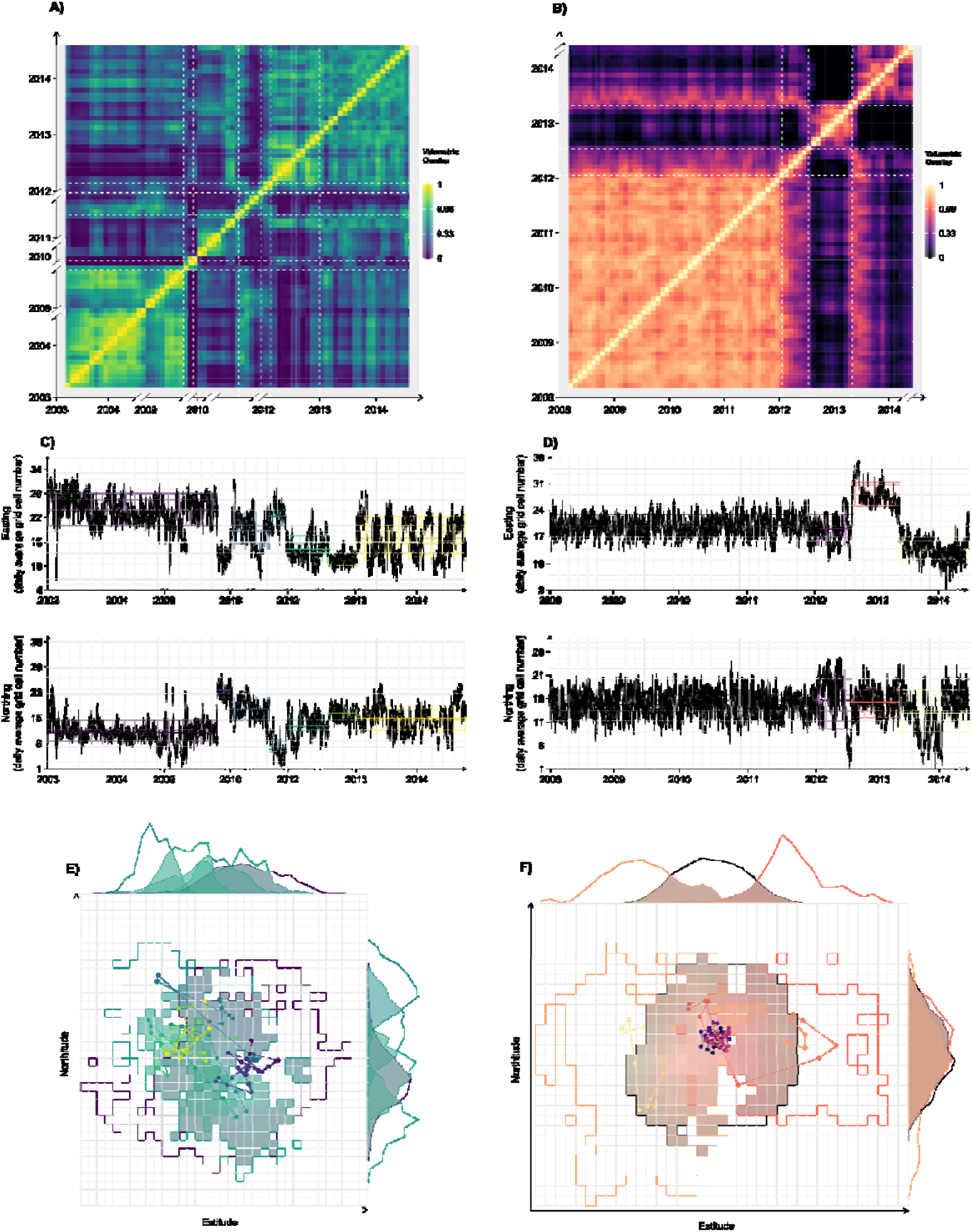
Shift occurrences and associated patterns based on the grid data set, which were available for CAR1 (left) and CAR2 (right). In (A) and (C), the label for the year “2011” is not displayed for readability. (A, B) Dissimilarity matrix of overlaps. Lighter colours indicate higher values. White dashed lines indicate the occurrence of significant spatial shifts identified by the segmentation procedure. Time gaps > 1 month are indicated by the axis break signs. (C,D) Locational time series. The different stationary phases are identified by the corresponding mean location (plain horizontal line) and associated standard deviation (bandwidth) of different colours. (E,F) For clarity, only new home ranges following a large spatial shift are indicated (i.e., distance between barycentres > 1.6 km, see Methods). (lightest colours for the most recent dates).

##### Space use fidelity

We estimated the Utilization Distribution (UD) for each stationary phase. With the *grid data set*, the UDs were obtained simply by counting the number of times the various quadrats were visited, and the home ranges were defined by the set of quadrats visited at least twice (corresponding to cumulative isopleths of 88% to 97%). With the *GPS data set*, UDs were obtained through Biased Random Bridge/Movement-based Kernel Density Estimation (BRB/MKDE; Benhamou 2011; software freely available at www.cefe.cnrs.fr/fr/recherche/bc/dpb/216-simon-benhamou). Any two successive relocations were bridged only if the delay between them was < 30 min. Furthermore, successive locations within 30 m of each other were averaged (as small apparent movements are more likely to reflect GPS noise than actual movements), and the minimum smoothing factor was set to 100 m. The home-range areas and core areas were then determined as the minimum areas encompassed within the 95% and 50% cumulative isopleths of the UD, respectively. We computed the overlap between all pairs of UDs of the same group (separately for the *grid* and *GPS data sets*), using Bhattacharyya (1946)’s coefficient (see Fieberg and Kochanny 2005), which ranges from 0 (no overlap) to 1 (identical UDs):

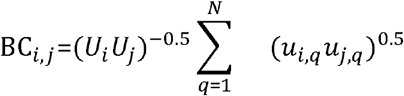

where *N* is the total number of quadrats involved (corresponding to the actual 250×250 m quadrats for the *grid data set*, and to virtual 10×10 m quadrats for the *GPS data set*), *u*_i,q_ and *u*_j,q_, are the fractions of UD encompassed in quadrat *q* for stationary phases *i* and j, and *U*_i_ and *U*_j_ are the total UD fractions encompassed in the home ranges (*U*_i_ = *U*_j_ = 0.95 for the *GPS data set*, while they took other, *ad-hoc* values in the range 0.88-0.97 for the *grid data set*). We then computed the spatial fidelity for each gorilla group (for the *grid* and *GPS data sets* separately) as the time-weighted average value:

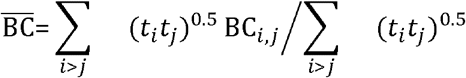

We considered that a large home range shift occurred whenever the distance between two successive home range barycentres was larger than the radius of the circle that has an area equal to the mean home range area (assessed from the *GPS data set*).

##### Recursion and Intensity distributions

For each stationary phase, in addition to the UD, we computed the Recursion Distribution (RD, i.e., the spatial distribution of revisits within the home range) and the Intensity Distribution (ID, i.e., the spatial distribution of the mean visit duration, Benhamou and Riotte-Lambert 2012, see Supplementary Material 4) using the BRB/MKDE program mentioned above. As RD and ID require a high spatial and temporal resolution to be computed, this was done only for the *GPS data set*. Any location was considered (re)visited when the resident group spent at least 12 min within 100 m of it. Given the gorilla mean travelling speed (about 0.4 km/h), the 12-min threshold corresponded to the minimum time required to fully cross the area. A new visit to this location was counted each time the group came back within 100 m of it after having left this area for at least 2 hours (the revisit to the area after a shorter delay was considered part of the same visit, and the additional time spent there was included in the duration of the visit). The 2-hour threshold corresponded to the minimum time required to ensure that the group really left the neighbouring of the location considered before coming back.

##### Daily path length (DPL)

We looked at how DPL, for each group *g* (among *G* groups) and day *d*, varied with the high and low frugivory season (*s*, dummy coded; data from transitory months between those two seasons (May and October; Masi and Breuer, 2018) were discarded), the group size (*z*), and the home range area (*A*) using a linear mixed Gaussian regression (Zuur et al. 2009):

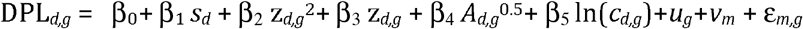

where β_0_ to β_4_ are the fixed effects, *c*_d,g_ is a control variable representing the tracking duration, and *u*_g_ ~ *N(*0, σ_g_) and *v*_m_ ~ *N(*0, σ_m_), with random effects σ_g_ *and* σ_m_, take into account the possible random differences between groups and months, respectively. The fixed and random effects are the parameters to be estimated by the statistical model, and ε_d,g_ is the error term. As years of study differed between groups, we could not include the year as a covariate. Details on statistical implementation are available in the *Statistical model implementation* section. We considered a quadratic effect of group size *z* to account for the multiple challenges of collective foraging (Papageorgiou and Farine, 2020). Indeed, larger groups should travel more in order to find more food. Yet, more group members may also slow down group movements (e.g., because of lower within-group coordination or of adaptation to the weakest members’ pace). Both these effects might be influenced by the group composition and, in particular, infants, as their energetic needs and ability to move drastically differ from adults. Thus, we fitted the model with and without 1-year-old (i.e., fully dependent) infants counted in the group size (as groups with more infants were not necessarily those with more adults, Supplementary Material Figure S7). We included the group identity in the possible factors influencing DPL to control for differences stemming from a ‘population’ effect and/or location sampling frequency (i.e., continuous vs. 20-min tracking in CAR vs. RC, respectively) and from group composition (e.g., number of adult or subadult males/females, adults/sub-adults/juveniles etc.). We considered it a fixed rather than a random factor to avoid assumption violation of residuals normality and homoscedasticity (see Supplementary Materials 6 and 7). We considered DPL values only for days for which gorillas were tracked for at least six hours (64% of the 1453 observation days). Furthermore, we filtered out potential upper outliers (DPL > 6 km), which represented about 1% of the remaining observation days, to minimise leverage effects that rendered the model less meaningful. The DPL distributions, before and after filtering, are presented in Supplementary Material Figure S4. For further details on model implementation, see *Statistical model implementation* below. As this analysis requires a high spatial and temporal resolution, we could conduct it only with the *GPS data set*.

##### Interactions between groups

We computed the two territoriality indices that are classically used in field primate studies: MR = 0.5 DPL/(A/π)^0.5^ (Mitani and Rodman 1979), and 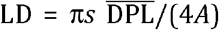 (Lowen and Dunbar 1994), where 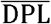 is the mean daily path length, *A* is the home-range area, and *s* is the mean ‘detection distance’. To further analyse the extent to which two groups with overlapping home ranges may have opportunities to interact with each other, we computed the UD overlap between neighbouring dyads, namely CAR1-CAR3 and RC1-RC2 (Figure 1), with the Bhattacharyya coefficient computed for periods during which the corresponding stationary phases overlapped in time. The time-weighted (to account for the length of the common duration of the used home ranges) mean overlap for a pair of neighbouring groups was then computed as

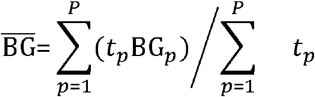

BG_p_ is the Bhattacharyya coefficient for a given period of time overlapping *p* (among *P* periods) whose duration is *t*_p_. We additionally computed the fraction of UD of one group encompassed in the area currently shared with the neighbouring group, FUD_1,2_ and FUD_2,1_ for any pair of neighbouring groups 1 and 2 (Smith and Dobson 1994; Benhamou et al. 2014). We then computed the time-weighted mean value 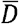 of the dissymmetry in space use, *D* = |FUD_1,2_ – FUD_2,1_| in the same way we computed 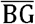. We also assessed the intensity with which a group may have used the central part of the home range of a neighbouring group. In the present study, the core area (i.e., the minimum area within the 50% cumulative isopleth of the UD) was highly fragmented, with many subparts occurring close to the home range periphery (Figure 1). We were therefore compelled to define the central part of the home range in another way. For this purpose, we defined the central part as the area within the 50% cumulative isopleths of the home range based on the elliptical home range model (Jennrich and Turner 1969; Mazurkiewicz 1969). It corresponds to a single elliptical area centred on the home range barycentre. We then calculated the number and average duration of visits of a group to the central part of its neighbour’s home range. A visit was counted as soon as the group entered this area. The number of visits was calculated for every month including at least ten tracking days. The average number of visits for each group was obtained using Poisson regressions. To determine whether the tendency of a group to enter the central part of a neighbouring group’s home range varied across the year, we fitted a Poisson model (see *Statistical model implementation*) *n*_m,g,k_ ~ Poisson *λ*_m,g,k_ with, *λ*_m,g,k_ = exp(β_0_+ β_1_ cos((*m*– 1+*k*)π/6)+*u*_g_+ln(*o*_m,g_)+ε_m,g_, where *n*_m,g,k_ is the number of visits for group *g* during the month *m* (January = 1; February = 2, etc.) and for a given temporal shift *k* (see below), *u*_g_ ~ *N(0*, σ_g_) where σ_g_ is the random effect on the intercept associated with the group identity and *o*_m,g_ is the associated offset term corresponding to number of monitored days that month for the group considered. We parametrised this model by the shift *k* to account for how *n*_m,g,k_ may vary across the year. The model was fitted to the data for each of the 12 possible *k* integer values in the range [0, 11], and we kept the *k* value for which we obtained the lowest Akaike Information Criterion (AIC, Burnham and Anderson 2002). Finally, we investigated the possible tendency that two neighbouring groups may have to move in close proximity, to avoid each other, or to independently move from each other when being simultaneously within a shared area. For this purpose, we computed the dynamic interaction index (Benhamou et al. 2014) using the “IAB” function of the *wildlifeDI* package (Long et al. 2014), between two groups *I*(*t*) = exp(−0.5(*d*(*t*)/c)^2^, where *d*(*t*) is the distance between the two groups at time *t*, and *c* is a critical threshold which smoothly separates large and small inter-individual distances. We tested whether the mean value of this index was significantly higher (attraction) or lower (repulsion) than expected under the null hypothesis of independent movements. The significance level is assessed through a time permutation procedure that maintains the actual order of locations (see Benhamou et al. 2014 for details). We considered three critical distances: 200, 500 and 1000 m, approximating critical distances expected for visual, olfactory and auditory cues in the forest environment based on the long-term experience we had observing gorillas at this site.

##### Statistical model implementation

Statistical models were fitted with either the *lme4* package (for mixed Gaussian models; Bates et al. 2015) or the *glmmTMB* package (for other models; Brooks et al. 2017). The singular effect of each variable was assessed by comparing model likelihood with and without the variable using a maximum likelihood ratio test with the “drop1” function, while, if multiple (DPL model), their synergic effect was assessed by comparing the *full* model, including all descriptive variables, with a *null* model, including only control variables. The ‘conditional’ variance explained by the model (i.e., including fixed and random effects) was calculated using the ‘r.squaredGLMM’ function from the *MuMIn* package (Barton 2020). To ease model interpretability, continuous control variables were centred and reduced (i.e., shifted and rescaled to have a null mean and unit variance, Schielzeth 2010). After fitting, we checked the necessary assumptions depending on the model family, using Q-Q plots, residuals vs. fitted values scatter-plots and histograms of residuals distribution (Zeileis and Hothorn 2002) based on the *DHARMa* package (Hartig 2022). The deviations obtained were reasonable (see Supplementary Material 6). We did not find overdispersion in the Poisson model used to investigate the number of visits. If several predictors were considered (‘DPL’ model), we also assessed the Variance Inflation Factor (VIF, Field 2005) using the ‘vif’ function of the *car* package (Fox and Weisberg 2019), which pinpointed several moderate correlation issues (max VIF > 3; “DPL” model) because some groups consistently had a higher weekly group size combined with low variance. The directions of average estimates were nonetheless robust: only significance was affected. For the ‘DPL’ model, when possible, we also checked model sensitivity to outlier points and random levels using Cook’s distance and dfBetas (Quinn and Keough 2003; Zuur et al. 2009) as implemented in the *influence*.*ME* package (Nieuwenhuis, Te Grotenhuis, and Pelzer 2012). This did not highlight major issues (Supplementary Material 7). We also considered the influence of the group identity by comparing model output (estimate and significance) with and without one group (Supplementary Material 7).

## RESULTS

### Site fidelity and space-use adjustment as a response to ecological and social challenges

All study groups except CAR3 shifted the location of their home range at some time(s) during the study periods. This occurred on average 0.72 time per year (range: [0, 1.8], Figure 3) sometimes to re-occupy an area previously used (e.g., CAR2; see Supplementary Material 3), but only 0.17 time per year when considering only large shifts (i.e > 1.6 km, corresponding to the radius of a circular area equal to the average home range size in our study). Despite they occupied multiple home ranges, these four study groups showed relatively high spatial fidelity (Bhattacharyya coefficients for *grid data set*: 0.36 and 0.39; for *GPS data set*: mean = 0.44, range = [0.24, 0.73]). The Simpson index was on average 0.43 for the *grid data set* (range: [0.36, 0.43]) and 0.51 (range: [0.25, 1]) for the *GPS data set*, indicating that, during the study period, gorillas spent most of their time exploiting a single area. The size of the home range (mean from all groups = 7.96 km^2^, range = [1.65, 13.48]; calculated from the *GPS data set* only) was positively correlated with the duration the group exploited this area (Spearman correlation: *r* = 0.56, *N* = 15, *P* = 0.03). Ten shifts out of 19 (*grid data set*: six out of nine, *GPS data set:* four out of ten) were larger than 1.6 km The *grid data set* provided further evidence that large spatial shifts occurred after long periods of stability and were often followed by periods of instability characterised by frequent small shifts (see CAR2 group after 2012, Figure 2) or by a drifting period with a gradual shift with no consistent spatial direction (see CAR1 group between 2010 and 2012; left panel, Figure 2). The across-season comparison showed that small (< 1.6 km) or large (> 1.6 km) rate of home range shifts was independent of the season (observed in high-fruit season: 0.16 (CI_95%_ = [0.03,0.4]) vs. expected: 0.33, *N* = 19, *P* = 0.14). We further looked at a plausible explanation for the various shifts made by three of the study groups for which we had associated behavioural data (CAR1, RC1, and RC2). Among the six large shifts done by CAR1 and RC1 (RC2 did only small-scale shifts), three were concomitant with the interaction with another gorilla group, another one occurred just after long heavy rain over several days, for which between-group encounters were more likely because of the difficulties of hearing other groups approaching, and another one occurred just after a visit to a distant aquatic area, where between-group encounters are also more likely because several groups share such areas. Eight out of nine of the small-scale shifts occurred just after (the same day or one or two days later) gorillas had an interaction with other gorillas (i.e., lone male or a group). All large and small shifts occurred far from the birth/arrival, or death/departure of group members (minimum absolute time distance of 35 and 28 days, respectively).

**Figure 3.**
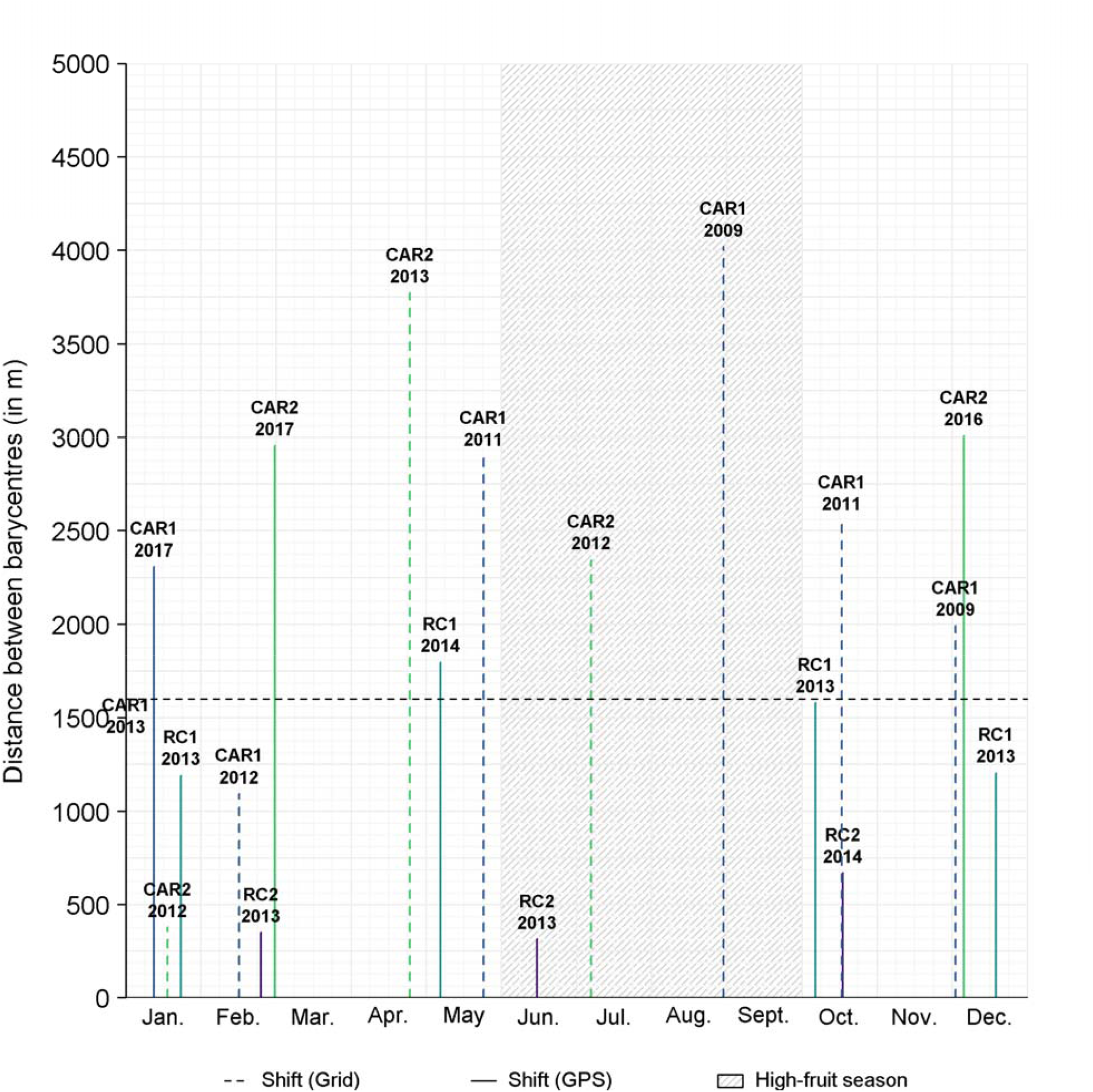
Small and large home range shifts| The period of high-fruit season indicates the period in which gorillas have a dominant frugivorous diet. The distance between the barycentres of two successive home ranges is indicated by the height of vertical bars. Plain and dashed vertical bars indicate values derived from the GPS and grid data sets. The name of each group is shown above the corresponding shifts. The dashed horizontal line indicates the threshold to distinguish between small and large home range shifts.

**Figure 4.**
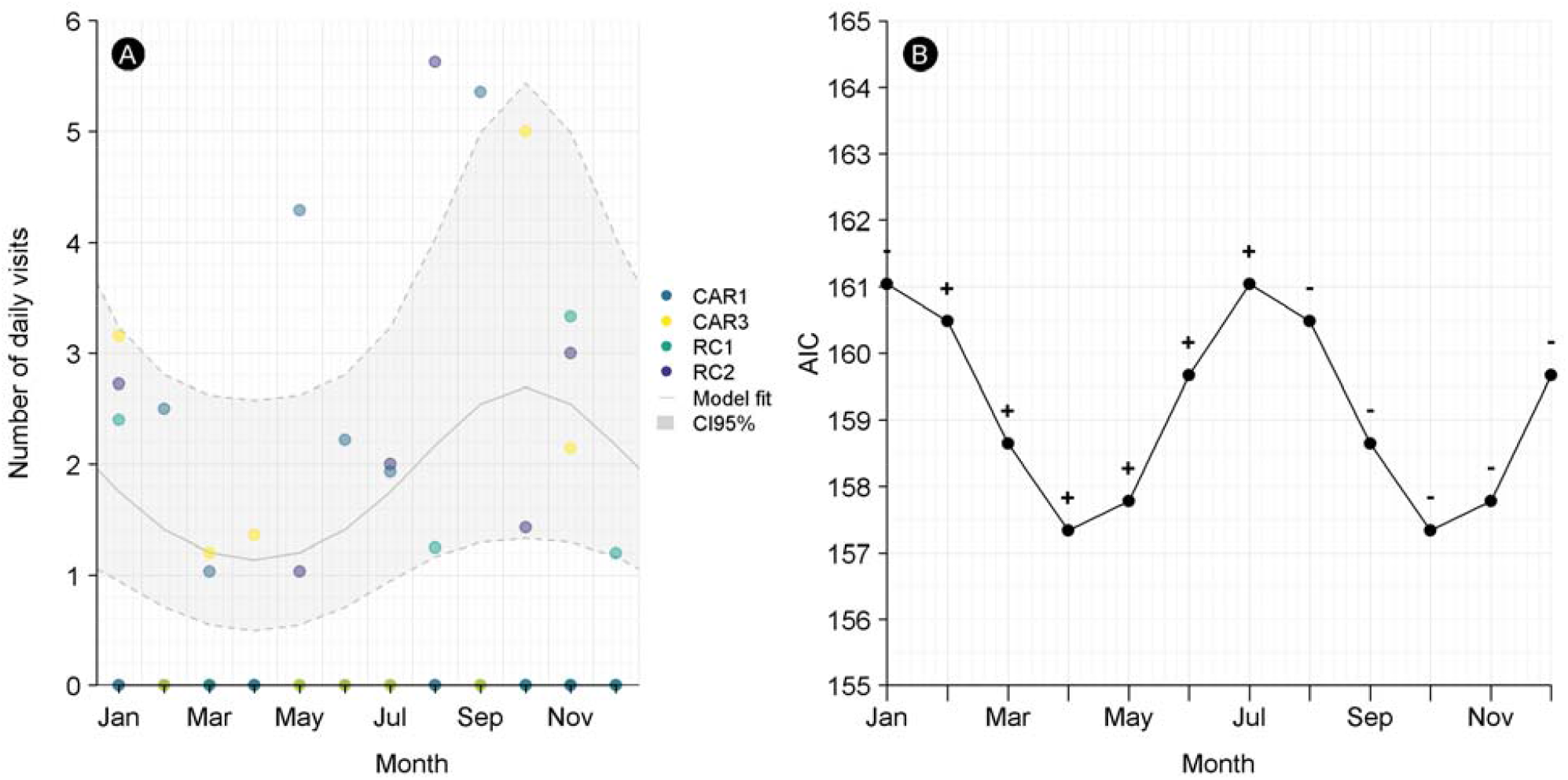
Number of daily visits to the central area of the neighbouring group’s home range | (A) Distribution of the number of daily visits for each month, year, and group. The best-fit model (based on the selection of the phase of the cosine, see B), holding the effect of other covariates/offset term at their average, is indicated by the plain grey line, while the associated 95% confidence interval is indicated by the grey background bordered with dashed lines. (B) AIC of preliminary generalised linear mixed models investigated seasonality in visit number within the central area of the neighbouring group’s home range as a function of the month on which the cosine variable was centered on (i.e., the phase). The model formula is shown in Methods, Group interactions, third paragraph. The model presented in (A) is the one for which the phase maximised the fit (i.e., minimised the AIC). The signs close to the dots indicate the sign of the estimated regression slope associated with the month.

### Uneven use of space within the home ranges

The UDs have a mean “flatness” (ratio between the size of the core area and the home range size depicting how evenly space is used) of 0.25 (CI_95%_ = [0.22, 0.28]), i.e., close to the flatness of a bivariate Gaussian distribution (ca. 0.23). However, the core area was not central but fragmented over the whole home range area (Figure 1). Overall, gorillas stayed at the most frequently/highly visited sites for the longest time at each revisit (Figures S2 and 3), independent of season (paired Wilcoxon signed rank test comparing the overlap between the ID and RD during high- and low-fruit seasons: *V* = 12, *P* = 0.461). When the same area was used over several years, sites highly and regularly used varied across seasons (larger overlap of the ID and RD within season compared with between season, Figure S2; see also Figure S3 for a visual representation of IDs/RDs).

### Season and group size constraints shape daily path length

The CAR groups seemed to travel more than the RC groups (Table 1). However, subsampling the CAR location data to match the RC protocol (a location recorded every 20 min as opposed to continuous collection of CAR data) resulted in a DPL estimate reduced by 638 m ± 162 m (see Supplementary Material Figure S6). This is close to the mean difference observed between the CAR and RC groups (DPL model with infants, Table 1, in line with McCann et al. 2021). Variation in DPL was significantly explained by some of the tested predictors (*full* vs. *null* model comparison: *χ*^2^ = 21.54, df = 4, *P* < 0.001; pseudo-*R*^2^ = 0.52). This variation was mainly driven by a seasonal effect, as gorillas travelled more during the high-fruit season (with low-fruit season taken as reference, est. = 79.55, CI_95%_ = [689.81, 747.84], *χ*^2^ = 21.07, df = 1, *P* = 0.002, Table 1, Figure 5). Home range size correlated poorly with DPL (*χ*^2^ = 1.73, df = 1, *P* = 0.19, Figure 5). The effect of group size was significant (*χ*^2^ = 8.16, df = 1, *P* = 0.004). This took the form of an inverted bell shape (negative estimate of squared terms, Table 1). This shape was flattened when < 1-year-old dependent infants were not considered within the group size (Figure 5, Supplementary Material Figure S6).

**Table 1.** Summary of the generalised linear regressions. Est. = Estimate, SE = Standard error, CI_95%_ =t he 95% confidence interval, Df = Degree of freedom, χ^2^ = Chi-squared statistics. Absent values or those with no meaning are not displayed (i.e., ‘-’ symbol). The level for categorical variables is indicated between parentheses. Scaled continuous variables are indicated as such, their initial mean and SD are depicted in footnotes. Duration of the day was scaled in the DPL model. Prior to transformation, mean = 8.46, SD = 0.95

| Variable | Est. | SE | CI<sub>95%</sub> | Df | $\chi^2$ | P |
| --- | --- | --- | --- | --- | --- | --- |
| Model Visit Core Area: (N = 71) |  |  |  |  |  |  |
| Intercept | -3.69 | 0.31 | [-4.29,-3.09] | - | - | - |
| cos(month number shifted at best) | 0.43 | 0.23 | [-0.01,0.87] | 1 | 3.76 | 0.052 |
| Model DPL (with infants):<br>(N = 1188) |  |  |  |  |  |  |
| Intercept | 254.25 | 655.65 | [-1014.56,1541.6] | - | - | - |
| (home range area)<sup>0.5</sup> | 79.55 | 60.44 | [-38.2,197.12] | 1 | 1.73 | 0.188 |
| Season (Frugivory) | **520.28** | **113.34** | **[302.39,739.16]** | **1** | **21.07** | **0.002** |
| Squared group size | **-22.25** | **7.79** | **[-37.34,-6.9]** | **1** | **8.16** | **0.004** |
| Group size | **378.9** | **140.43** | **[102.24,651.07]** | **1** | **7.28** | **0.007** |
| Log of tracking duration | **175.58** | **25.74** | **[125.08,225.66]** | **1** | **46.54** | **<0.001** |
| Group name (CAR2) | **194.69** | **122** | **[-45.17,431.64]** | **4** | **76.97** | **<0.001** |
| Group name (CAR3) | **1321.89** | **106.34** | **[1113.76,1529.16]** |  |  |  |
| Group name (RC1) | **-435.52** | **249.23** | **[-924.91,49.19]** |  |  |  |
| Group name (RC2) | **-837.73** | **112.25** | **[-1056.57,-618.16]** |  |  |  |
| Model DPL (without infants):<br>(N = 1188) |  |  |  |  |  |  |
| Intercept | 1412.96 | 472.63 | [501.53,2341.34] | - | - | - |
| (home range area)<sup>0.5</sup> | 47.6 | 63.48 | [-75.6,171.55] | 1 | 0.56 | 0.454 |
| Season (Frugivory) | **504.68** | **113.16** | **[287.15,723.21]** | **1** | **19.89** | **0.002** |
| Squared group size | -4.6 | 6.29 | [-16.76,7.81] | 1 | 0.54 | 0.464 |
| Group size | 109.75 | 103.48 | [-95.07,309.57] | 1 | 1.12 | 0.289 |
| Log of tracking duration | **169.43** | **26.15** | **[118.1,220.3]** | **1** | **41.98** | **<0.001** |
| Group name (CAR2) | **111.85** | **119.64** | **[-122.29,345.13]** | **4** | **74.49** | **<0.001** |
| Group name (CAR3) | **1342.21** | **99.75** | **[1146.96,1536.62]** |  |  |  |
| Group name (RC1) | -<br>**1136.16** | **249.81** | **[-1623.85,-647.76]** |  |  |  |
| Group name (RC2) | **-972.35** | **117.63** | **[-1200.17,-740.45]** |  |  |  |

**Figure 5.**
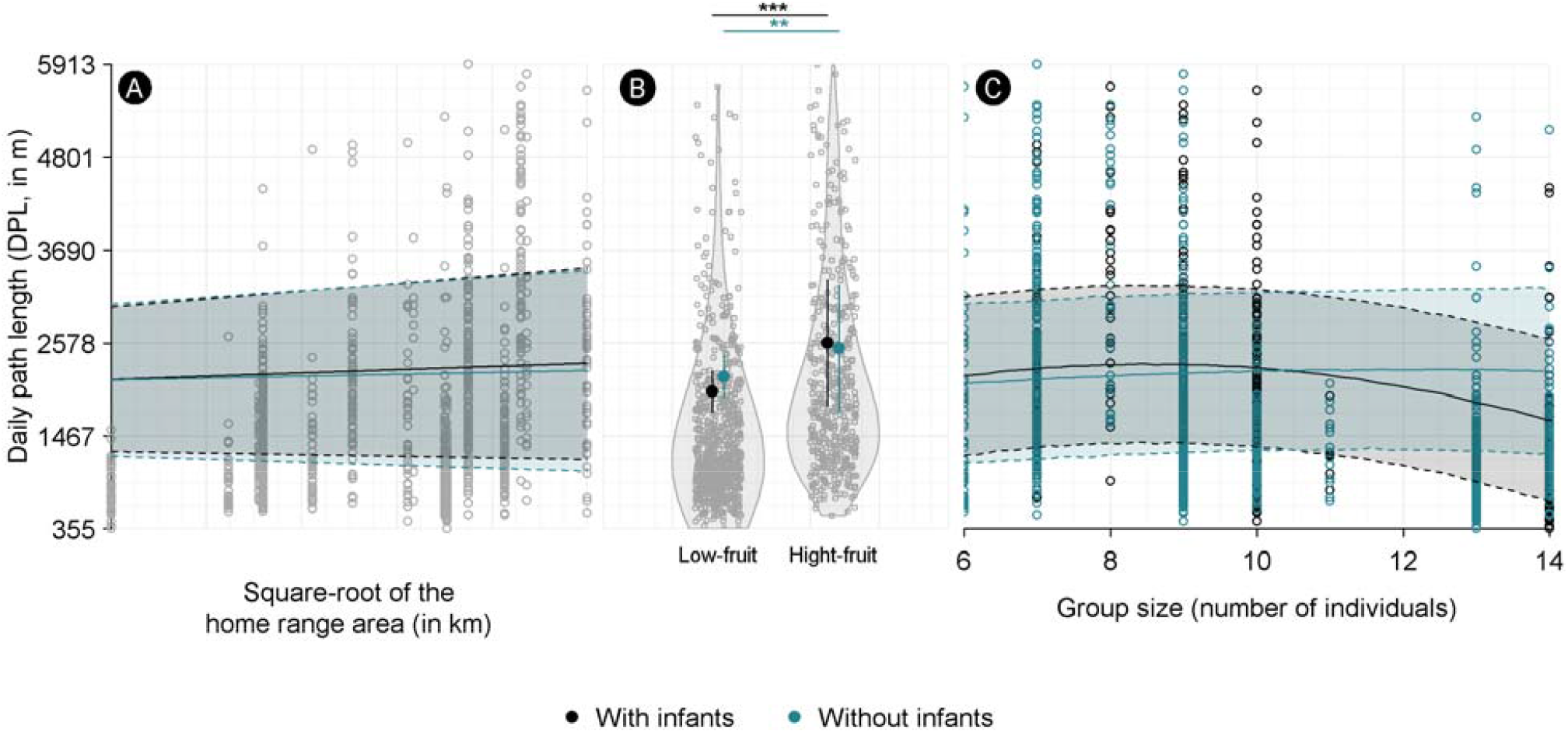
Effect of home range size (A), dietary season (B) or group size (C) on daily path length | As the effect of one predictor is conditional on other predictor values, we plotted the predictor of interest’s effect when the effect of any other predictor was averaged. For the group identity, we did so by averaging predictions for each group. The result is indicated by the plain line. The confidence interval is plotted with a coloured background for continuous variables or with vertical bars for categorical variables. The models presented here either include less than 1-year-old infants (black and white colours), or not (blue). Similarly, raw data are indicated by points, with grey points. In B), the background density estimation of data distribution (‘violin’ plot) is either prior (black) or after (coloured) dependent infants have been removed. In C), confidence intervals were obtained using the ‘predictInterval’ function of the *merTools* package (Knowles and Frederick 2020).

### Inter-group spatial relationships suggest absence of territoriality

The two territoriality indices were low (MR: mean = 0.49, CI_95%_ = [0.41, 0.56], LD: mean = 0.08, CI_95%_ = [0.07,0.10]); Supplementary Material Figure S1), far below the thresholds (1 and 0.08) empirically assumed to distinguish between non-territorial and territorial species and were highly correlated (Spearman correlation: *N* = 15, *r* = 0.75, *P* < 0.002). The mean UD overlap (Bhattacharyya coefficient) between neighbouring groups was very low in both dyads (CAR: mean = 0.08, range = [0, 0.2]; RC: mean = 0.04, range = [0, 0.19]; Table 2). The level of overlap was unrelated to the time duration that dyad groups used this area (Spearman correlation: r_s_ = −0.49, *N* = 10, *P* = 0.15). For each dyad, the shared area was used as intensively by both dyads of neighbouring groups (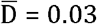 for both dyads, with values close to 0 indicating symmetry in use of shared space by both groups within a dyad; Table 2). A group ventured into the central part of the neighbouring group’s home range on average 0.59 times a month (range= [0.31, 1.58], see Supplementary Material Figure S1 for group-level value), with a maximum frequency just before the start of the high-fruit season (April; lowest AIC for the model including for k = 4;*χ*^2^ = 3.76, df = 1, *P* = 0.05, Supplementary Material Figure 5). Within a dyad, no group visited the central area of the other group more often than the other dyad group (paired Wilcoxon signed rank test with continuity correction: CAR dyad, *V* = 23, *P* = 0.68, RC dyad, *V* = 22, *P* = 0.21). The mean duration of these visits was 115 min (CI_95%_ = [46, 29]). In two visits (out of 41), the group stayed even 12 h within the central area of its neighbouring group, spending the night there. Within the RC dyad, neither group visited the central part of the home range of the other dyad group for longer time than its neighbour did (Welch’s t-test: *N* = 16, *t* = 0.8, df = 13.71, *P* = 0.44). Within the CAR dyad, the CAR1 group visited the central part of the home range of CAR3 for a significantly longer time than the CAR3 visited the CAR1 central area (mean difference of 262.5 min, CI_95%_ = [28.9, 496.1], t-test: *N* = 25, *t* = 2.39, df = 15.08, *P* = 0.03; Residuals were normally and homogeneously distributed, see Supplementary Material 8). On average for both dyads, the two neighbouring groups were within 500 m of each other only 5.11 times a year. At their closest, the two groups were observed at 150 m (RC dyad) and 288 m (CAR dyad) of each other. Overall, groups tended to move independently of each other (permutation test of locations, most *P* > 0.05, Table 1). Only attraction was significantly observed twice (CAR dyad, parameter threshold > 500 m; but not at a lower threshold, Table 2).

**Table 2.** Comparison of observed and expected values of dynamic interaction levels using a randomisation procedure. Comparisons were computed for RC and CAR dyad separately for each period during which home-range overlapped (periods of overlap with few simultaneous tracks within the overlapping area were discarded). The *p*-values correspond to a one-tailed test. Observed interaction levels significantly lower and larger than the expected values means avoidance and joint movements, respectively. The *c* values correspond to the distance threshold considered.

| Critical distance | Synchronized relocation number | Interaction (observed) | Interaction (expected) | *p*-value |
| --- | --- | --- | --- | --- |
| **RC dyad (1st common period)** |  |  |  |  |
| *c* = 200 m | 510 | 0.153 | 0.204 | 0.361 |
| *c* = 500 m | 510 | 0.538 | 0.632 | 0.206 |
| *c* = 1000 m | 510 | 0.826 | 0.876 | 0.09 |
| **RC dyad (2nd common period)** |  |  |  |  |
| *c* = 200 m | 333 | 0.016 | 0.024 | 0.372 |
| *c* = 500 m | 333 | 0.26 | 0.255 | 0.402 |
| *c* = 1000 m | 333 | 0.552 | 0.533 | 0.468 |
| **CAR dyad** |  |  |  |  |
| *c* = 200 m | 708 | 0.052 | 0.048 | 0.477 |
| ***c* = 500 m** | **708** | **0.395** | **0.255** | **0.01** |
| ***c* = 1000 m** | **708** | **0.748** | **0.599** | **0.001** |

## DISCUSSION

In this study, we provided a rare integrated longitudinal picture of the dynamics of space use by western gorillas at three temporal scales: everyday movements, home range behaviour (from several weeks to months) and ‘migrations’ (home range shifts, occurring a few times in several years). We focused on the effects of ecological (food and energy constraints, i.e., frugivory and group size, respectively) and social factors (group composition and neighbouring groups). We showed that the diet choice had a major influence on everyday movements, whereas social factors, such as intergroup interactions (thus potential for infanticide risks and lethal attacks; e.g., Watts, 1989; Potel et al. 2023), shaped movement patterns at the largest scales, leading to home range shifts. No territorial behaviour or space dominance were detected between neighbouring groups.

### Spatial shifts and socio-ecological factors

Overall, gorilla home ranges were generally less than 13 km^2^, while gorillas moved approximately 1.42 km per day (median value). Gorilla groups occasionally shifted their home range location, either locally (< 1.6 km, so that the new home range largely overlapped with the previous one) or more widely, except for the group tracked for the shortest time (1.5 years). Note however that this group has lately also shifted its home range of several dozen kilometres (unpublished Bai Hokou long-term data). However, sometimes, groups returned to an area previously used. This emphasises that western gorillas are relatively faithful to a given ranging area, as also found elsewhere (Scarry et al. 2023). Space use fidelity appears to be a ubiquitous trait in primates (Janmaat et al. 2009; Ramos-Fernandez et al. 2013; Campos et al. 2014; Wartmann, Juárez, and Fernandez-Duque 2014; Cheyne et al. 2019; Belle and Estrada 2020; José-Domínguez, Savini, and Asensio 2015; Caillaud et al. 2014). This is probably because of the advantages of greater familiarity with the environment, which is particularly useful for animals with high cognitive capacities to feed on spatio-temporally heterogeneous foods and/or to avoid predation (Powell and Mitchell 2012). This may be also particularly relevant when the group needs to maintain group cohesion and coordination, as it is the case for this species (Potel et al. 2023, Nellissen et al. 2024, 2026).

The onset of large shifts was followed by periods of instability in the space use, involving frequent small shifts (Figure 2A and B), presumably to readapt to new ecological (e.g., food distribution/availability) and social (e.g., presence of other gorilla groups/solitary males) conditions. Such a behaviour was also observed in mangabeys that become less cohesive, move more, and venture into larger areas per day when settling into a new area, likely to gain information on the new environment (Janmaat and Chancellor 2010). Although the activity budget, diet, and movement patterns of western gorillas are strongly influenced by seasonal changes in fruit availability (Remis 1997; Goldsmith 1999; Cipolletta 2004; Doran-Sheehy et al. 2004; Masi, Cipolletta, and Robbins 2009; Masi et al. 2015), the rate of home range shifts (small and large) was independent of the season. These shifts occurred irregularly, leading to high variability in the lifetime of the home ranges (i.e., for how long a given area was used). However, although we could have also missed some auditory interactions of the focal group with other gorilla groups or with elephants, our behavioural observations suggest that large shifts might be related rather to social interactions. By contrast, small shifts may be mainly related to male reproductive strategies, as previously observed for a former group of western gorillas in CAR, which moved its home range to acquire females before returning to its original range after a sub-adult female joined the group (Cipolletta 2004). In western gorillas, females are known to disperse when they reach sexual maturity (Stokes, Parnell, and Olejniczak 2003; Masi et al. 2021). Males (solitary or in groups) could therefore travel long distances to try to find potential dispersing females from other groups (Masi et al, 2021) or, on the contrary, to avoid other groups’ home range to prevent their own females from dispersing (Breuer, Robbins, and Robbins 2016; Cipolletta 2004). Although we did not observe shifts related to female acquisition (or loss), the vast majority of small shifts occurred after interacting with other social units, risky events in which extra-group males may lead to infanticide, death of the silverback and/or female transfer (Stokes, Parnell, and Olejniczak 2003; Baudouin et al. 2019; Manguette et al. 2020; Potel et al, 2023).

### Ecological factors shape everyday movements

Movement decisions should be based on current and/or future environmental conditions, as well as metabolic and nutritional needs (Nathan et al. 2008), which may vary across seasons and group sizes. We found that these two factors (but not the home range size considered as a proxy of defensibility) affected the daily path length (DPL). This is consistent with previous evidence that dietary changes shape gorilla activity and travel time or distances (Remis 1997; Goldsmith 1999; Cipolletta 2004; Doran-Sheehy et al. 2004; Masi, Cipolletta, and Robbins 2009; Seiler and Robbins, 2020). The effect of dietary season was larger when compared to the positive effect of group size (Figure 5). Indeed, finding sufficient food for maintaining nutritional and energetic balance in a seasonal environment is a crucial nutritional challenge for western gorillas (Masi et al. 2015; Masi and Breuer, 2018), particularly for large groups (Fuh et al. 2022). They travelled more in the high-fruit season than in the low-fruit season, echoing previous observations (Remis 1997; Goldsmith 1999; Cipolletta 2004; Doran-Sheehy et al. 2004; Masi, Cipolletta, and Robbins 2009; Seiler and Robbins 2020). In the low-fruit season, western gorillas mostly feed on herbs and leaves (Masi et al. 2009, 2015), which are usually permanently available, homogeneously distributed and occurring at high density, as opposed to fruit (Milton 1981; Janmaat et al. 2016). This suggests that more than the energy requirements of the group and the energy contained in the targeted food, the variation in movement patterns might be the result of variation in the challenge of locating resources such as fruit (Robira et al. 2023a).

However, nutritional challenges and food choice may also depend on group size (Fuh et al. 2022). When fully dependent infants (< 1-year-old) were included, the group size had a significant effect on DPL, which did not appear when they were excluded, with larger groups moving shorter DPL than smaller groups did (Figure 5 and Table 1). In addition to the infanticide risk forcing the group sometimes to travel more to avoid conspecifics, the presence of infants is likely associated with a higher energy demand for lactating mothers (Masi, Cipolletta, and Robbins 2009; Lodwick and Salmi 2019). This should lead to a larger DPL to find more food. Yet, infants may also contribute to slowing down the group, which should lead to shorter DPL. The observed relationship highlights this trade-off. This is why groups with an intermediate size may forage more efficiently, as frequently reported in literature (Markham et al. 2015; Papageorgiou and Farine 2020). As the pattern disappeared when group size did not include infants, this unlikely results from a group inertia stemming from accrued coordination. This result supports instead the idea that the movement of western gorilla groups is framed around the weakest group members, such as infants, likely to ensure the crucial group cohesion. Comparatively, group size (without non-weaned individuals) U-shaped the distances travelled in mountain gorillas (Grueter et al. 2018). This is supposedly explained by considering that inter-group competition is a preponderant constraint on foraging efficiency, and is considerably lowered the larger the group. This constraint may appear because of their increased population size, which has recently led to closer proximity, with an exacerbation of aggressive behaviour between neighbouring groups (Caillaud et al. 2020). As we emphasise in the next paragraph, such direct competition appears weak in western gorillas, which should explain why the U-shape pattern is unrelated to adult-only group sizes. This comparison exemplifies how major influence of local constraints (compared to evolutionary constraints) can shape space-use behaviour, hence possibly major differences between phylogenetically close species.

### Passive spatial segregation

The benefits of self-restraining space use to a limited area, in terms of knowledge about the current food distribution, tend to decrease when space is shared with conspecifics because they are likely to deplete the same resources (Powell and Mitchell 2012, Riotte-Lambert et al. 2015). We showed that western gorillas tended to use exclusive space. This resulted in little overlap between home ranges of the neighbouring groups considered in our analyses (which likely represent only a small subset of the actual neighbouring groups/solitary males). This raises questions about the driving factors of this spatial segregation: direct aggressive behaviour (i.e., territoriality) or indirect passive segregation (Riotte-Lambert et al. 2015). Based on data obtained with camera traps on non-habituated groups, Morrison et al. (2020) suggested that western gorillas are territorial because the areas most used by one group (i.e., core areas) were avoided by others. In our study, we highlighted that habituated groups ventured into the central part of the neighbouring habituated group’s home range on average more than once every two months. Furthermore, these visits could be long (median duration of visits: 156 min), and a group may sometimes stay overnight there (two out of 41 visits). Although the central part of the home ranges may not correspond to the core area, our result showed at least that a group could venture far inside the home range of a neighbouring group. The low overlap we found between neighbouring home ranges agrees with Morrison et al. (2020)’s results, but we claim that the avoidance of areas intensively used by another group does not necessarily involve territoriality. Many groups of western gorillas frequently visit the same distant large swamp clearings (Magliocca, Querouil, and Gautier-Hion 1999), sometimes located very far away from the centre of their home range (Robira et al 2024). Therefore, they would certainly need to cross other groups’ home ranges to reach them. Altogether, these results underline that, despite space being barely shared with neighbouring groups, gorillas are not fierce defenders of space. It suggests that they do not consider places further away from their own home range centre as riskier than others. Accordingly, we did not observe patrols, and the DPL was lower than expected for a territorial animal that would attempt to defend an area of an equivalent size. However, it is worth noting that territoriality indices, which combine DPL and home range area, should be considered with caution because the values of both DPL and home range area dramatically depend on the methods used to measure them, making comparisons across studies difficult to interpret. Furthermore, we did not detect any large asymmetric relationships between the neighbouring groups, as would be expected in the case of the existence of direct competition. The spatial behaviour of the neighbouring groups in both study dyads was very similar despite the different group sizes (used here as a proxy for group strength): groups used the home range of the neighbouring group relatively equally, exploited the same locations in shared areas equally intensively, and ventured into the neighbour’s home range equally frequently, and for equal periods of time. In contrast, larger groups of mountain gorillas had exclusive use of more than 50% of their home range, while smaller groups had less than 20% (Caillaud et al. 2014). Additionally, groups did not actively avoid each other, suggesting that general avoidance was likely due to passive mechanisms.

Like other Hominidae, western gorillas are known to rely on spatio-temporal long-term memory to forage (Trapanese et al. 2019, Salmi et al. 2020; Robira et al. 2023a, b; Robira et al. 2024). In our study, the sites exploited for the lengthiest times were also the most visited. They changed seasonally (Supplementary Material 4). These observations converge on the prevalence of long-term spatial memory shaping foraging decisions. It is therefore likely that the memorisation of depleted areas by others contributes to the observed pattern of passive spatial segregation (i.e., without territoriality, Riotte-Lambert et al. 2015). Potentially, long-range multimodal signalling, such as with vocalisations or odour (Salmi and Doran-Sheehy 2014; Miglietta et al, 2020; Nellissen et al., 2026) should also promote fast identification of known opponents thus long-range passive avoidance (i.e., avoiding direct fights; Masi and Bouret 2015).

## CONCLUSION

This study highlighted how food may affect directly (by changes in spatio-temporal distribution) and indirectly (because of inter-group scramble competition), the daily and seasonal space use strategy of frugivorous species inhabiting a seasonal environment. As fruit can be limited during certain periods of the year (Janmaat et al. 2016; Masi et al. 2015), this may have been relevant for shaping the species’ spatiotemporal cognitive abilities (Milton, 1981; Rosati, 2017; Ashton et al. 2020). The space use strategy, and the cognitive level supporting it, might not only be shaped by the competition with conspecifics, but also many other sympatric species (Havmøller et al. 2021; Robira and Perez-Lamarque, 2023). Promising avenues hence lie in considering multi-species dynamics to better understand the patterns of species segregation and space use particularly for aiding the conservation strategies of endangered species.

## ETHICS

All research protocols adhered to the ethics rules of the local country at time of the study where it was conducted (Central African Republic and Republic of the Congo). Research permits were provided by the Ministries in Central African Republic and Republic of the Congo under the support of World Wide Fund for Nature (WWF) and Wildlife Conservation Society (WCS), respectively.

## DATA AVAILABILITY

Data will be accessible upon reasonable requests to TB for the RC data set, and to ShM and TF for the CAR data set. Analytic codes will be provided by BR upon request afterwards.

## ACKNOWLEDGEMENTS

The work was conducted under the permission of the “Ministère de l’Economie Forestière et du Développement Durable” as well as the “Ministère de la Recherche Scientifique et de l’Innovation Technologique” of the Republic of Congo, as well as the “Ministères de l’Education Nationale et de l’Enseignement Technique, et des Eaux et Forêts, Chasse et Pêche” of Central African Republic (CAR). We deeply thank the Wildlife Conservation Society and their associated teams, as well as local field assistants and the Aka trackers for tremendous involvement in data collection. We thank Emmanuel Konyal for help to gather social data. We also deeply thank the team of the Primate Habituation Program of Dzanga-Sangha Protected Areas (DSPA) and the Administration of DSPA and WWF CAR, as well as the invaluable Aka trackers from Bai-Hokou and Mongambe for their help and contribution to this work. We thank the USAID Central Africa Regional Program for the Environment (CARPE) and Spain-UNEP LifeWeb for financially supporting Mondika Research Center. We thank the Action Transversal of the Museum National d’Histoire Naturelle (MNHN), the Projet Federateur of the Human and Environment Department of the MNHN and the UMR 7206 and Cleveland Metropark Zoo for their financial support for the CAR study. This study is part of the long-term Studies in Ecology and Evolution (SEE-Life) program of the CNRS. BR was funded by a PhD scholarship from the French Ministry of Higher Education and Research to the “École Normale Supérieure de Paris.” We thank Emiel van Loon and Karline Janmaat for discussion, and Luigi Boitani for his support. We thank Ran Nathan and two anonymous reviewers for constructive feedback on previous versions of this manuscript.

## AUTHORS’ CONTRIBUTION

BR conceived the study with SB and ShM, BR cleaned and analysed the data, and wrote the first draft. ASM, AT, CC, EOB, GB, SiM, ShM, and TF collected the data. SB, ShM and TB primarily revised the article. SB and ShM subsequently revised it several times. All authors revised and finally approved the final version.

