## Supplementary Material for "Spatial segregation and multi-scale space-use adjustments to socio-ecological variables in western gorillas"

### 1 Group composition

A detailed composition of groups during the GPS study period is shown in Table S1. Exact group composition during the grid data period is not available.

Table S1: Group composition at times of the GPS study. Age/sex class was established based on Breuer et al. [2009]. SB = Silverback, AF = Adult female, SAM = Sub-adult male, JUV = Juvenile, INF = Infant (M = Male, F = Female). Absence of data of first and last observation indicates that gorillas were present before and after the study period. Dates follow the year/month/day format.

| Locality | Group | ID | Mother | Age/Sex class | First observation | Last observation |
|---|---|---|---|---|---|---|
| CAR | 1 | A | - | SB | - | - |
| CAR | 1 | B | - | AF | - | - |
| CAR | 1 | C | - | AF Votre texte ici 1 | - | - |
| CAR | 1 | D | B | SAM | - | - |
| CAR | 1 | E | - | JUV | - | - |
| CAR | 1 | F | B | INF | - | - |
| CAR | 1 | G | B | INF | - | - |
| CAR | 2 | H | - | SB | - | - |
| CAR | 2 | I | - | AF | - | - |
| CAR | 2 | J | - | AF | - | - |
| CAR | 2 | K | - | BB | - | - |
| CAR | 2 | L | - | SAM | - | - |
| CAR | 2 | M | - | JUV | - | - |
| CAR | 2 | N | I | JUV | - | - |
| CAR | 2 | O | J | JUV | - | - |
| CAR | 2 | P | I | INF | - | - |
| CAR | 2 | Q | J | INF | 2016-11-01 | - |
| CAR | 3 | R | - | SB | - | - |
| CAR | 3 | S | - | AF | - | - |
| CAR | 3 | T | - | AF | - | - |
| CAR | 3 | U | - | AF | - | - |
| CAR | 3 | V | - | AF | - | - |
| CAR | 3 | W | - | SAM | - | 2017-03-01 |
| CAR | 3 | X | T | JUV | - | - |
| CAR | 3 | Y | V | JUV | 2016-07-01 | - |
| CAR | 3 | Z | U | JUV | - | - |
| RC | 1 | AA | - | SB | - | - |
| RC | 1 | BB | - | AF | - | - |
| RC | 1 | CC | BB | SAM | - | - |
| RC | 1 | DD | BB | INFM | - | - |

Table S1: Group composition at times of the GPS study. Age/sex class was established based on Breuer et al. [2009]. SB = Silverback, AF = Adult female, SAM = Sub-adult male, JUV = Juvenile, INF = Infant (M = Male, F = Female). Absence of data of first and last observation indicates that gorillas were present before and after the study period. Dates follow the year/month/day format. *(continued)*

| Locality | Group | ID | Mother | Age/Sex class | First observation | Last observation |
|---|---|---|---|---|---|---|
| RC | 1 | EE | - | AF | - | - |
| RC | 1 | FF | EE | SAM | - | - |
| RC | 1 | GG | EE | INFM | - | - |
| RC | 1 | HH | - | AF | - | - |
| RC | 1 | Q | HH | JUVF | - | - |
| RC | 1 | JJ | - | AF | - | - |
| RC | 1 | KK | JJ | INFM | - | - |
| RC | 1 | LL | - | AF | - | - |
| RC | 1 | MM | LL | INF | 2013-05-18 | - |
| RC | 1 | NN | - | Y-AF | - | - |
| RC | 2 | OO | - | SB | - | - |
| RC | 2 | PP | - | AF | - | - |
| RC | 2 | QQ | - | INFM | 2013-01-21 | 2014-06-07 |
| RC | 2 | RR | - | AF | - | - |
| RC | 2 | SS | - | SAF | - | 2014-08-12 |
| RC | 2 | TT | - | JUVM | - | - |
| RC | 2 | UU | - | Y-SB | - | - |
| RC | 2 | VV | - | Y-SB | - | 2014-07-20 |
| RC | 2 | WW | - | SAF | - | 2014-08-12 |
| RC | 2 | XX | - | BB | - | - |
| RC | 2 | YY | - | Y-AF | - | 2012-12-09 |
| RC | 2 | ZZ | - | Y-AF | - | 2013-01-28 |

#### 2 Raw data visual summary

We report in Figure 1 the raw estimates for each group of the several space use descriptors which were used for the analyses.

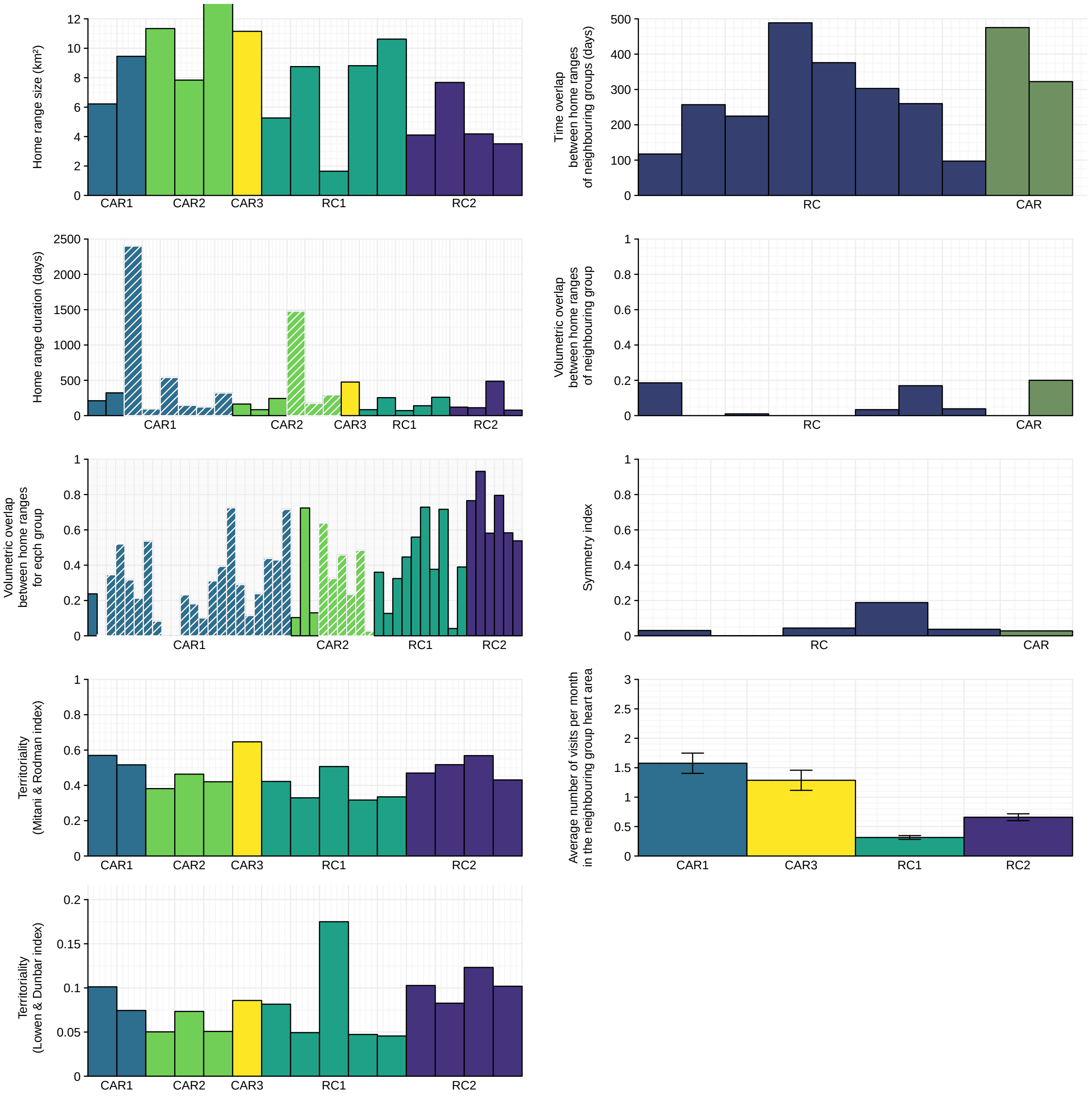


Figure S1: Visual summary of calculated indices associated to intra-group space-use pattern (left) or inter-group space-use pattern (right) | Indices are presented within group for each home range, or each pair of home ranges for inter-group indices. In this latter case, at the notable exception of the general overlap, home range pairs when no overlap occurred were obviously discarded. Each group is evidenced by its own colour matching colours throughout all figures. Dyads are identified by the mixing of respective groups' colour. The number of vertical bars differs between groups/dyads as a consequence of the differences in number of home ranges used during the study period. Grid data are emphasized by the dashed pattern. The vertical segments indicate the standard error.

### 3 Segmentation procedure

We present below the output(s) of the bivariate segmentation [“segmentation” function of the *segclust2* package, Patin et al., 2020], on the northing and easting coordinates of the group (GPS data), so as to determine stationary locational series for the study groups. For each group, the daily locations (and the associated rough paths) are presented in the top panel, and coloured according to the stationary phase, identified by the segmentation procedure, to which they belong. The time series of daily locations $(x,y)$ is shown below. The different stationary phases are identified by their mean (plain horizontal line) and their standard deviation (bandwidth) of different colours.

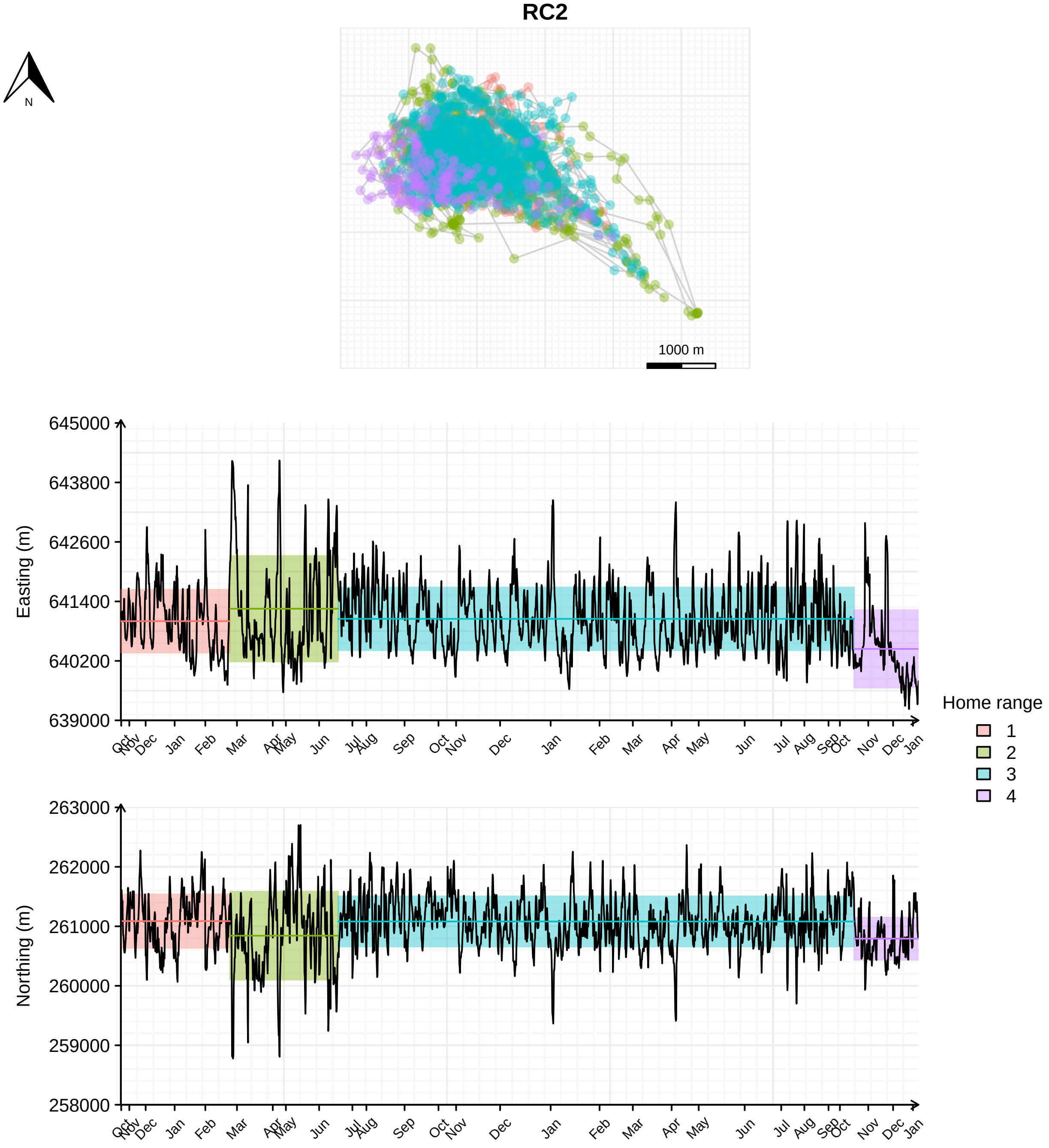

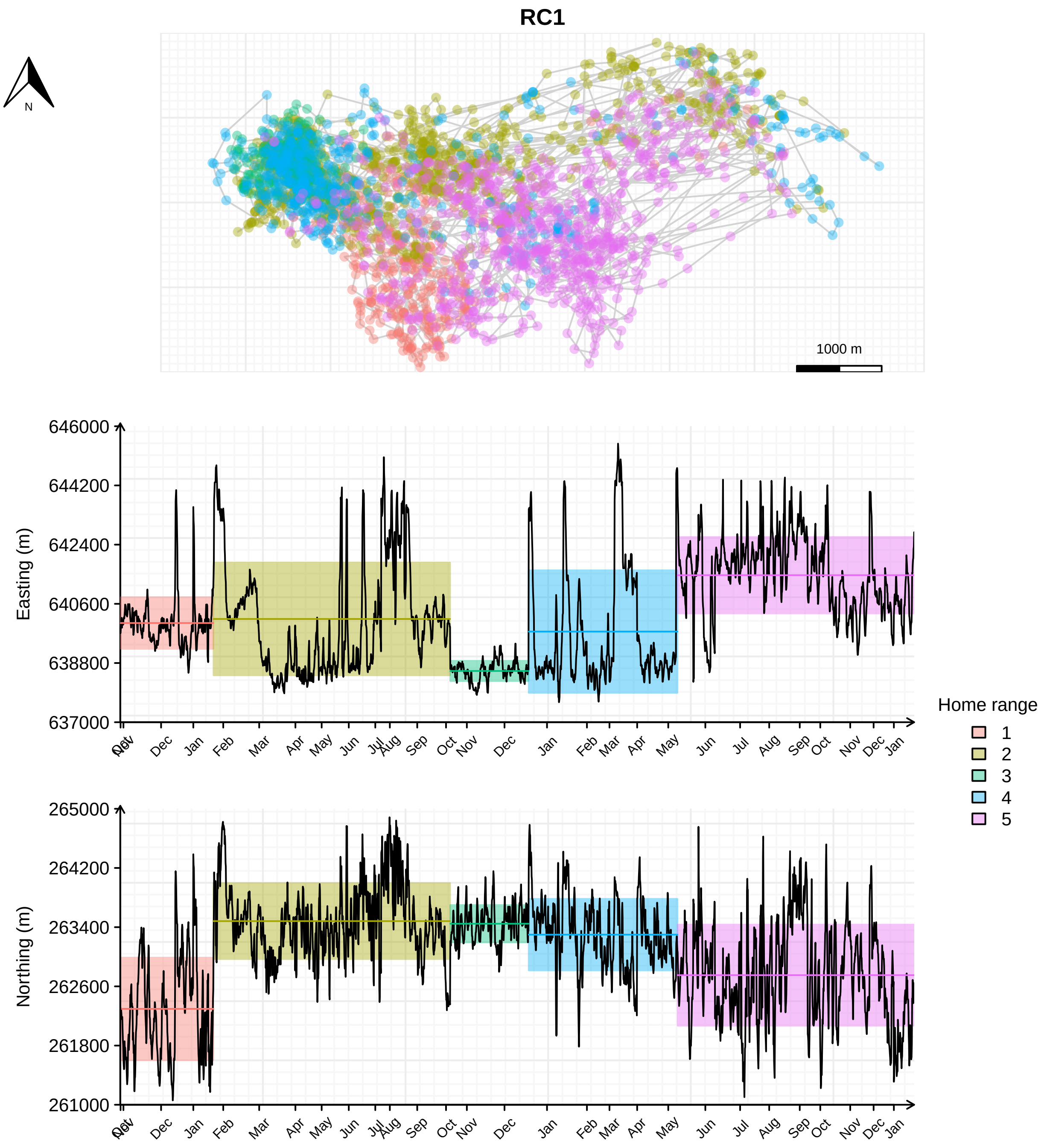
RC1
N
1000 m
646000
644200
642400
640600
638800
637000
Easting (m)
Oct
Nov
Dec
Jan
Feb
Mar
Apr
May
Jun
Jul
Aug
Sep
Oct
Nov
Dec
Jan
Feb
Mar
Apr
May
Jun
Jul
Aug
Sep
Oct
Nov
Dec
Jan
265000
264200
263400
262600
261800
261000
Northing (m)
Home range
1
2
3
4
5

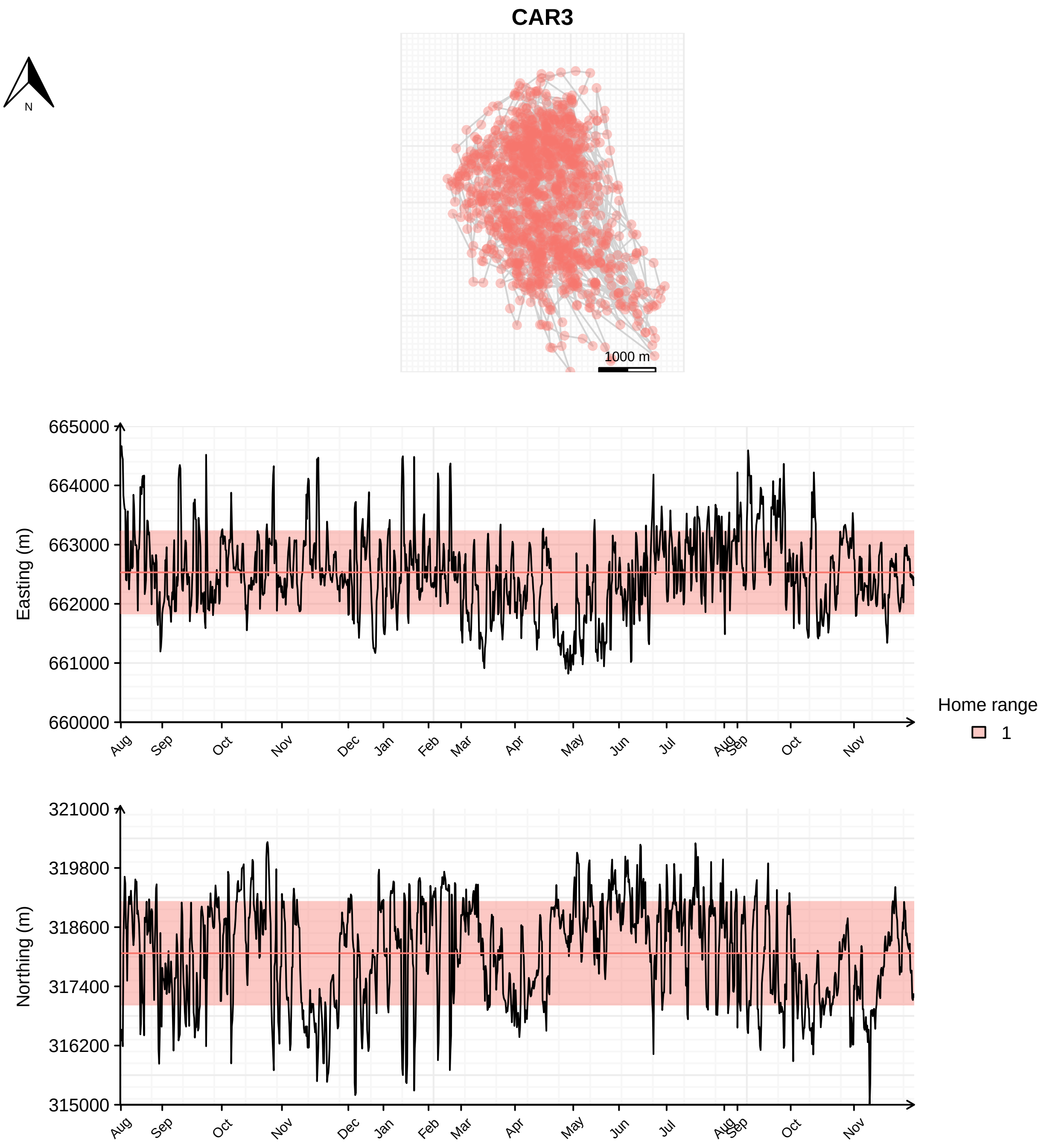
CAR3
N
1000 m
Easting (m)
665000
664000
663000
662000
661000
660000
Aug
Sep
Oct
Nov
Dec
Jan
Feb
Mar
Apr
May
Jun
Jul
Aug
Sep
Oct
Nov
Home range
1
Northing (m)
321000
319800
318600
317400
316200
315000

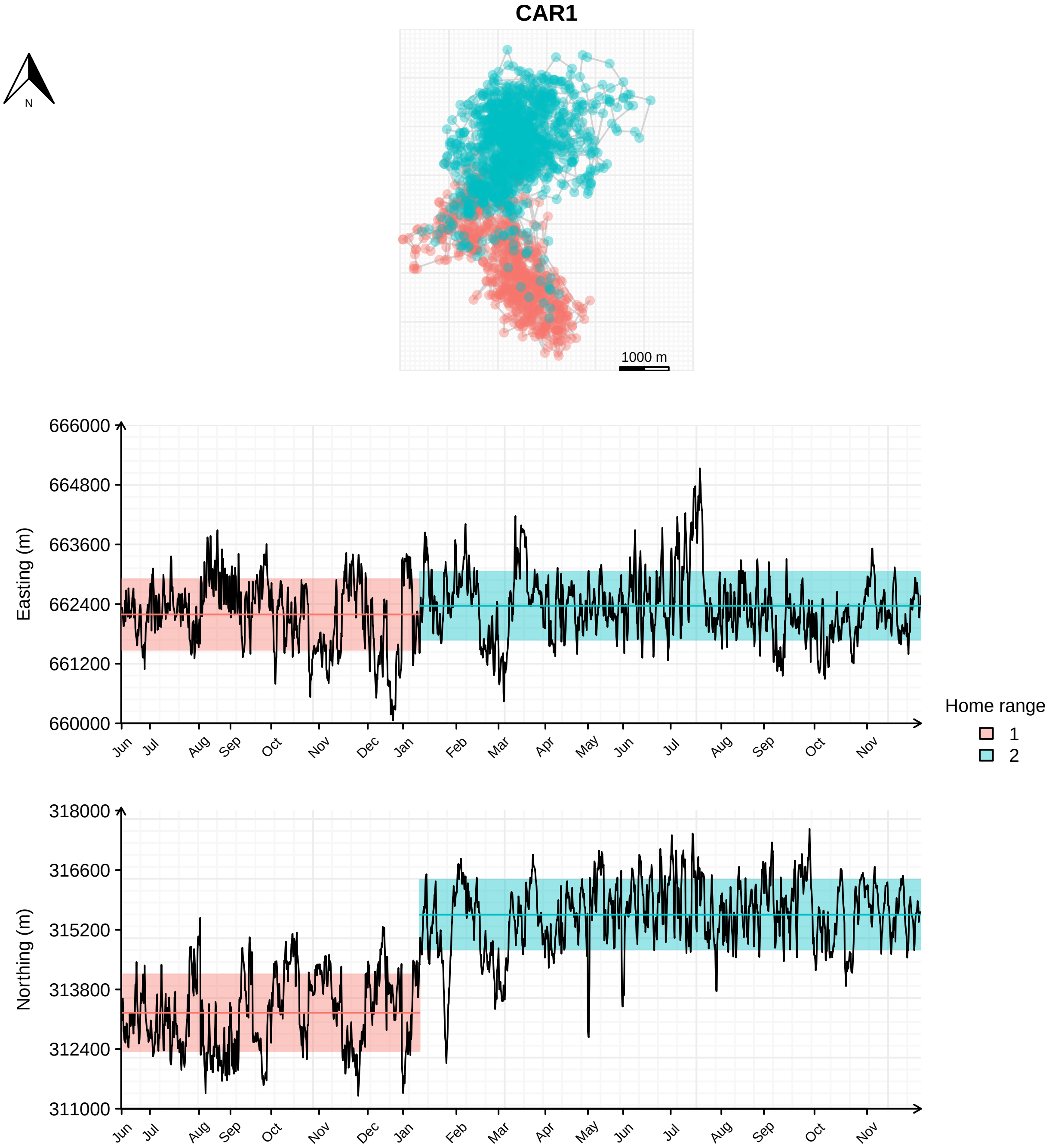

CAR1
N
1000 m
Easting (m)
666000
664800
663600
662400
661200
660000
Jun Jul Aug Sep Oct Nov Dec Jan Feb Mar Apr May Jun Jul Aug Sep Oct Nov
Home range
1
2
Northing (m)
318000
316600
315200
313800
312400
311000
Jun Jul Aug Sep Oct Nov Dec Jan Feb Mar Apr May Jun Jul Aug Sep Oct Nov

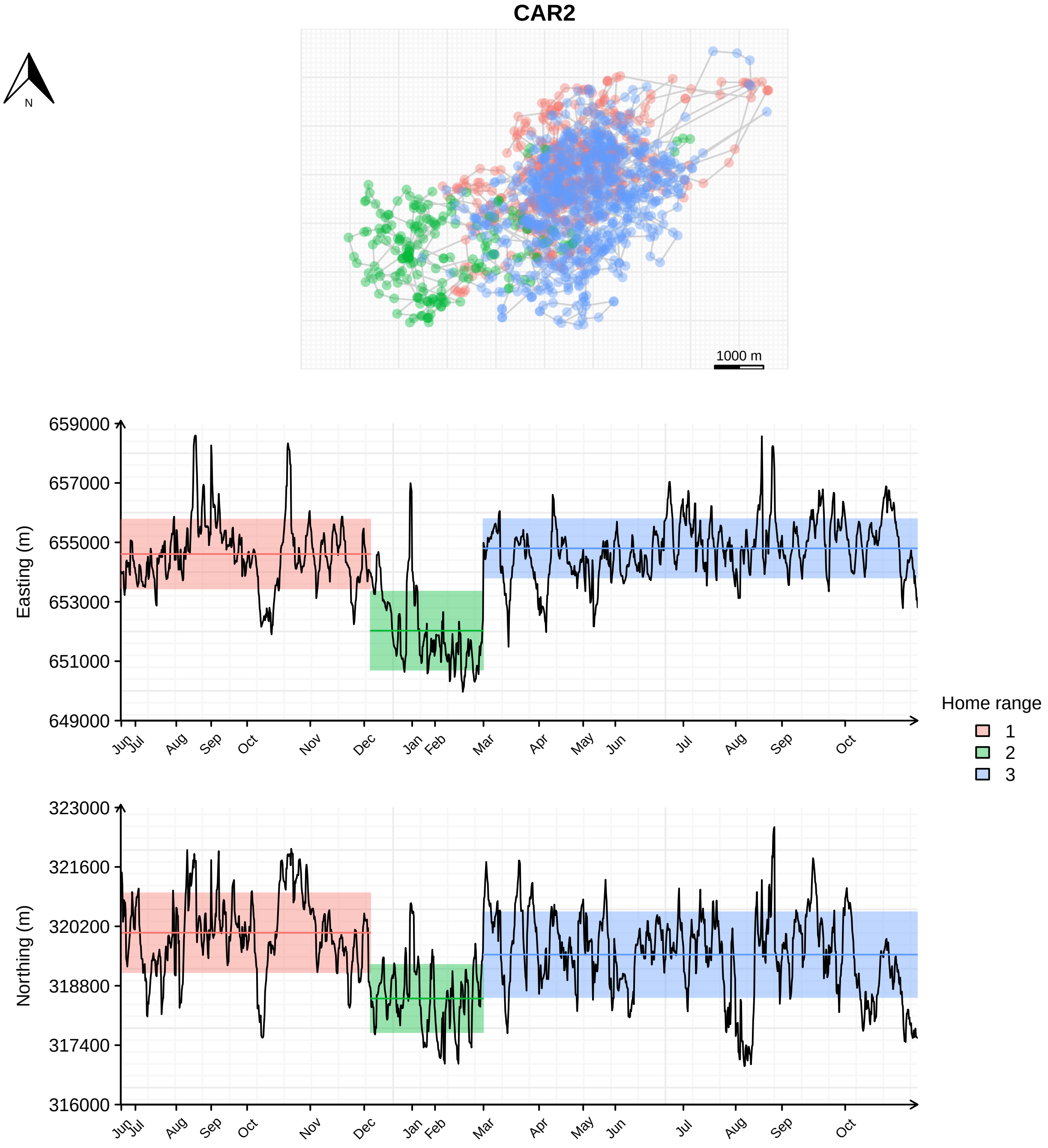
CAR2
N
1000 m
Easting (m)
659000
657000
655000
653000
651000
649000
Northing (m)
323000
321600
320200
318800
317400
316000
Jun Jul Aug Sep Oct Nov Dec Jan Feb Mar Apr May Jun Jul Aug Sep Oct
Home range
1
2
3

### 4 Intensity Distribution (ID) and Recursion Distribution (RD) overlap

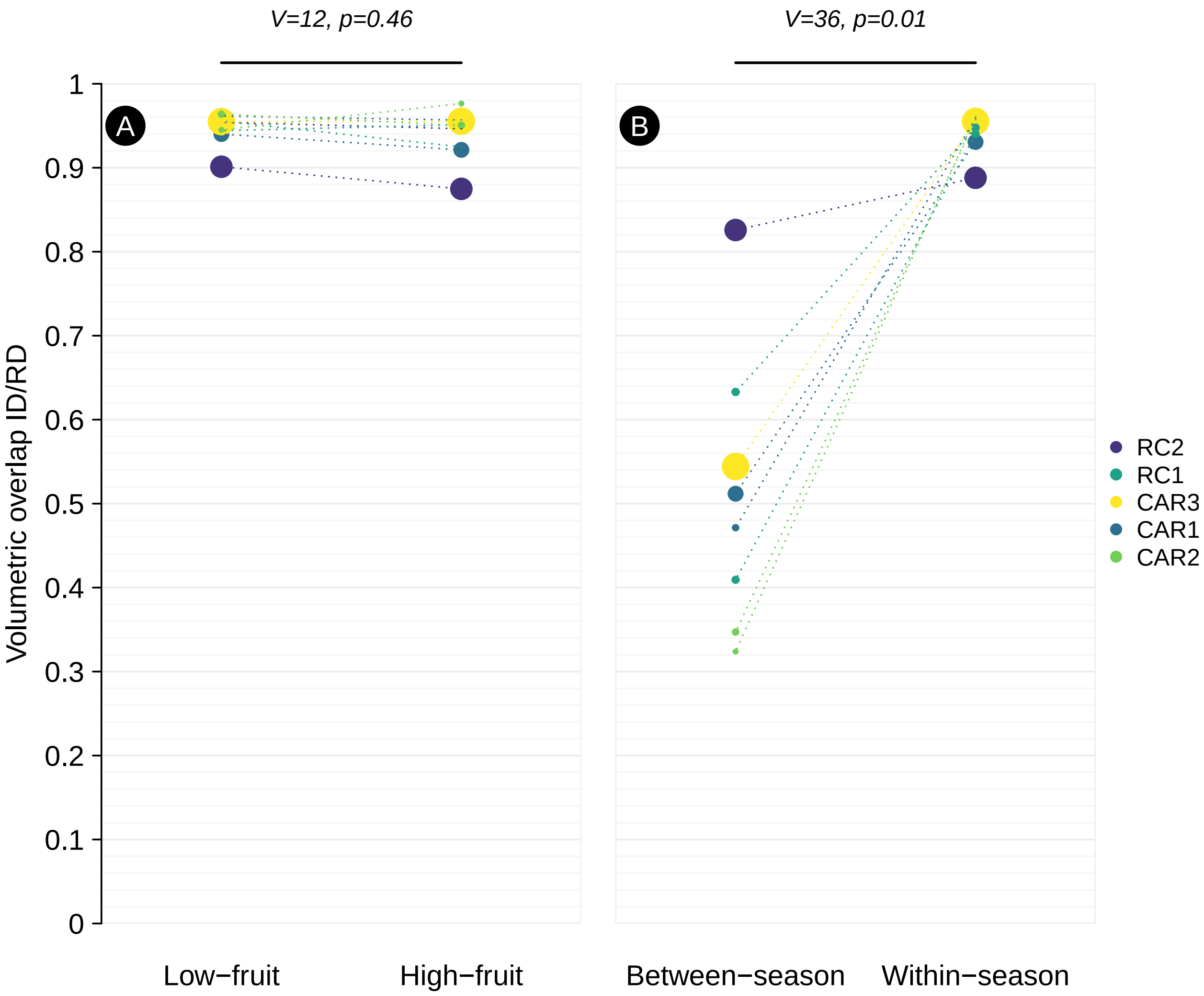


Figure S2: Volumetric overlap between the Intensity Distribution (ID) and Recursion Distribution (RD) (A) during the high-fruit (May-September) and low-fruit (November-March) season (B) within a same season or between the two seasons | Only home ranges including the high-fruit and low-fruit season are indicated. Paired data are linked by the dashed lines and were compared using a paired Wilcoxon signed rank exact test ("wilcox.test" function). The sizes of the circles are proportional to the home range duration.

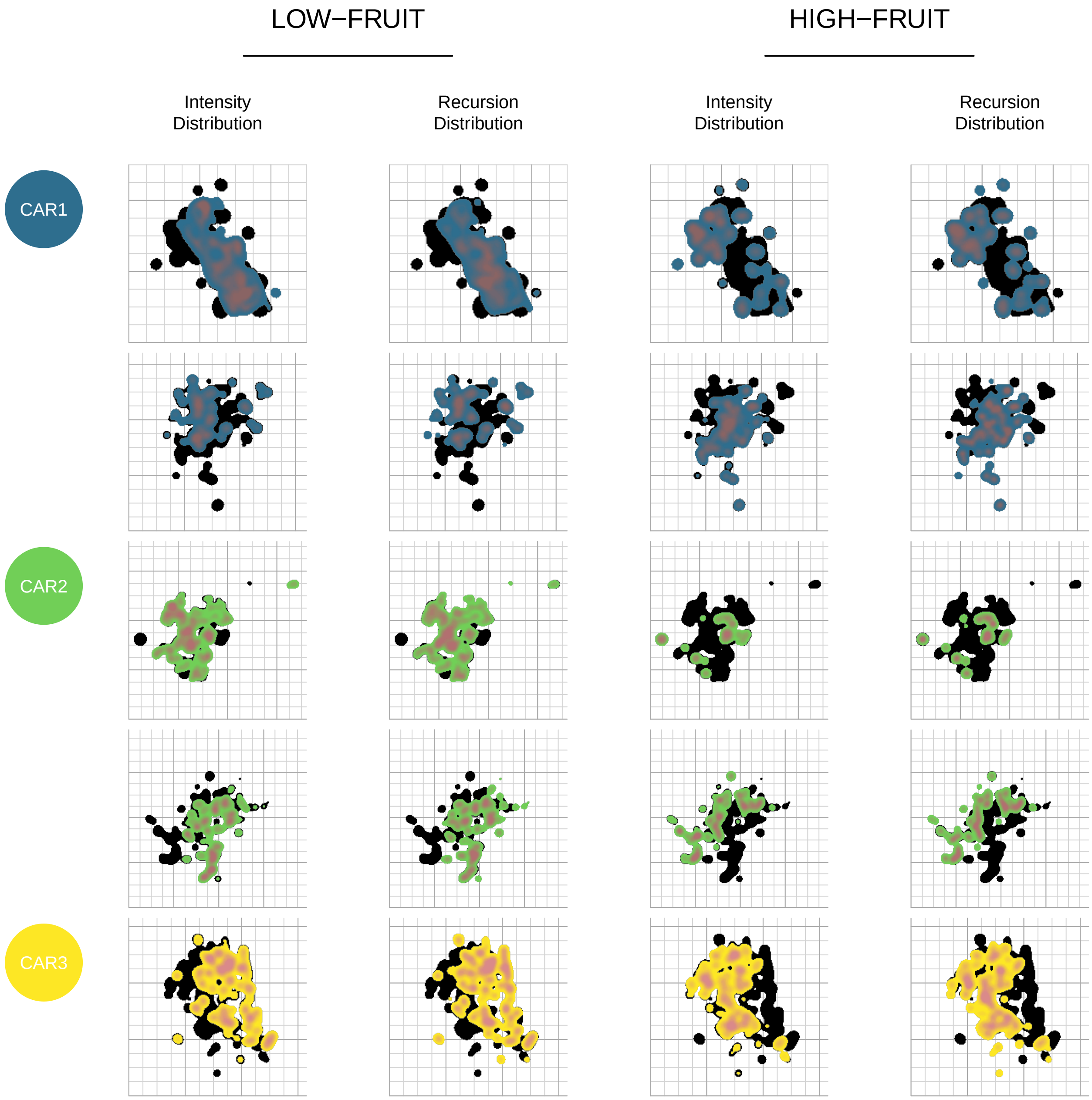


Figure S3: Intensity Distribution (ID) and Recursion Distribution (RD) for each CAR group, home range and season | Colours match those of each group in other graphics. The colour intensity highlights the value of the distribution (darker if the relative value is higher). The black background shows the whole home range. The background grid shows cells of 500 m of side (lightgrey) or cells of 2000 m of side (darkgrey)

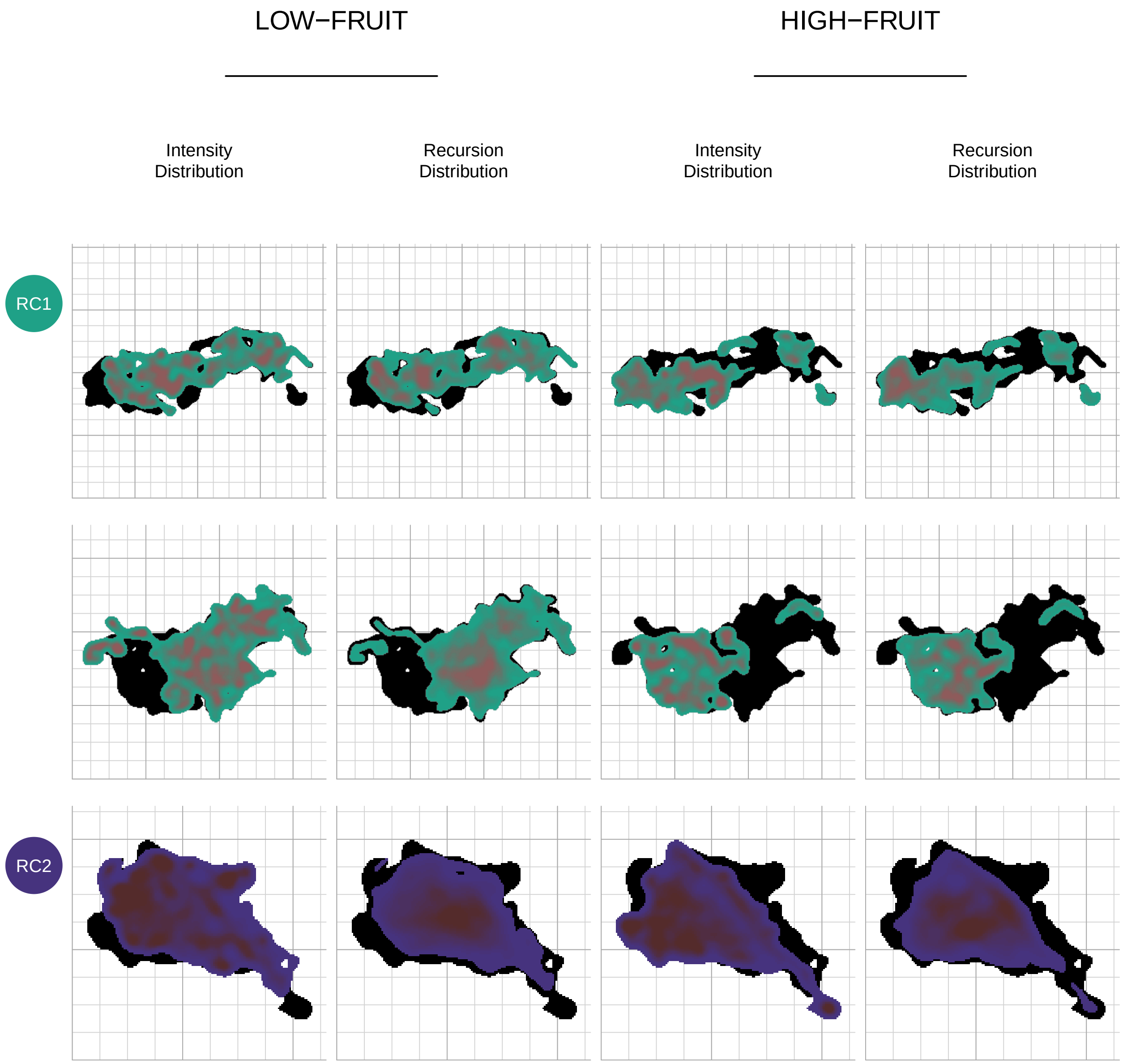
LOW−FRUIT
HIGH−FRUIT
Intensity Distribution
Recursion Distribution
Intensity Distribution
Recursion Distribution
RC1
RC2

#### 5 Daily Path Length (DPL) model

##### 5.1 DPL calculation

DPL was calculated for each day and group using the “SpatialLines” structure [*sp* package, Pebesma and Bivand, 2005, Bivand et al., 2013] and the “lineLength” function [*SDraw* package, McDonald and McDonald, 2020]. For continuous data, because the GPS can show erratic behavior, for example by recording close-distant points whilst the observer remains still (e.g. during feeding events), we re-discretized paths at a constant step length of 30 m to remove GPS noise. This was made possible using a home-implemented function relying on the *Rcpp* package [Eddelbuettel and François, 2011, Eddelbuettel, 2013, Eddelbuettel and Balamuta, 2018]. We did so only for this calculation, and not for home range estimation since the BRB/MKDE program (see Utilization Distribution (UD) estimation parameterization) already has an argument for filtering out inactivity. For subsequent analyses on DPL, we then filtered out DPL associated to (1) days with less than 6 hours of monitoring and (2) if they were of more than 6000 meters because they stand as extreme outliers, destabilizing the model estimation. A comparison of the distributions prior to and after filtration is shown in Figure S4.

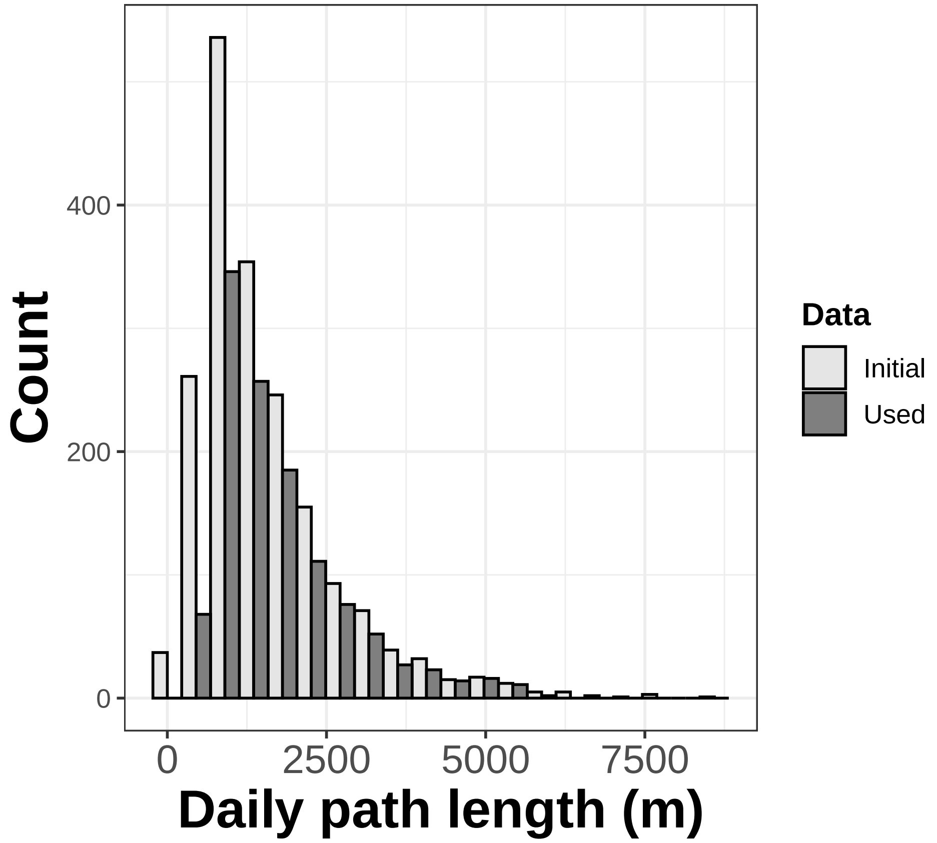


Figure S4: Distribution of the daily path lengtprior (initial) and after (used) filtration for days with insufficient monitoring ($< 6$ h), or unusually large values ($> 6000$ m).

#### 5.2 Influence of the sampling rate on DPL

We compared the DPL calculated with CAR data when considering **continuous** track-log or with a regular **20-min** sampling (as for RC data). This emphasizes that a sampling every 20 min reduces DPL estimation of 638 m (± 162) (Figure S5)

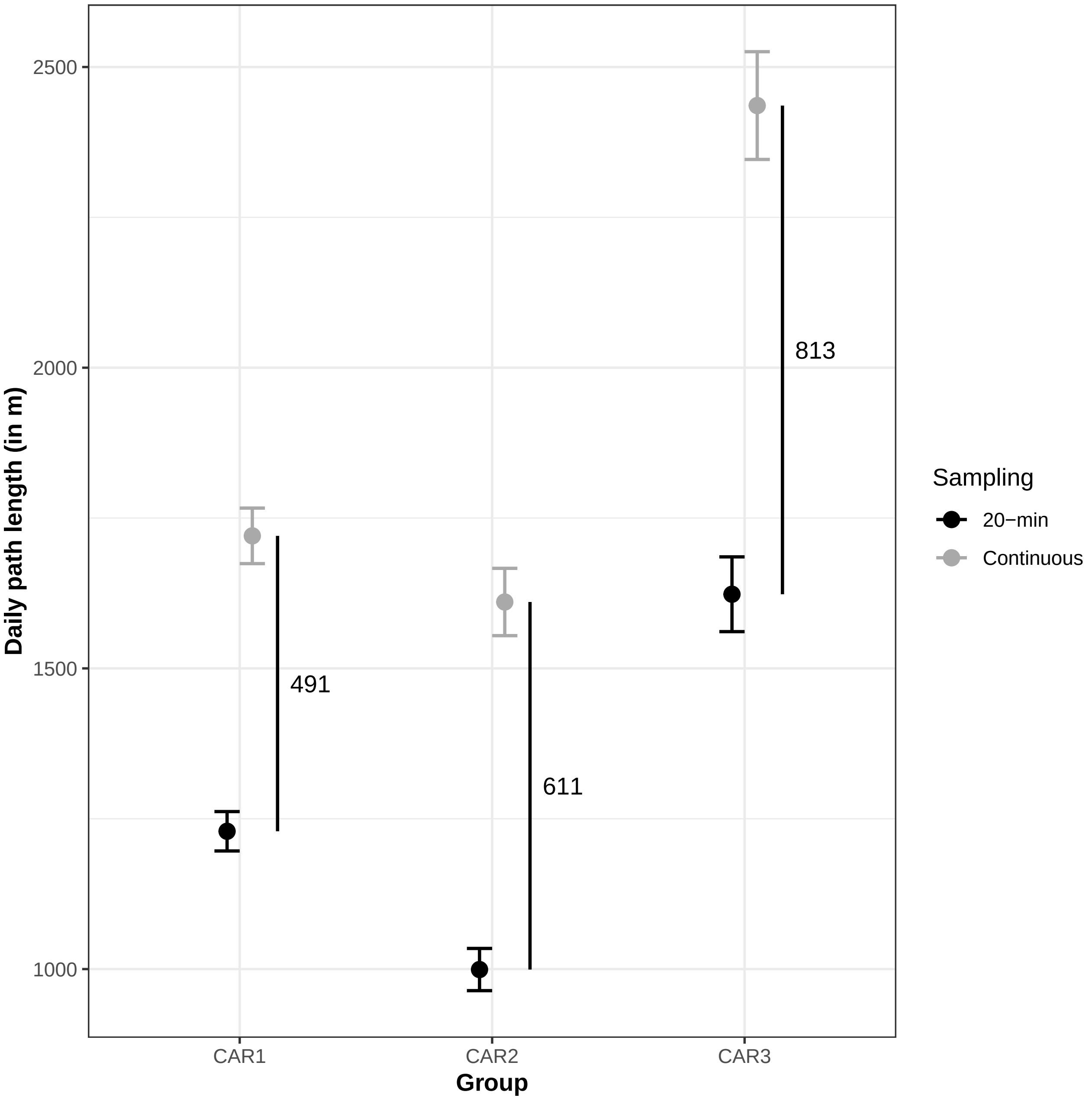


Figure S5: Location sampling protocole affects DPL estimation

#### 5.3 Discussion of the rationale for the DPL model implementation

We present below visual inspections of (generalized) linear (mixed) model assumptions, as well as various statistical indicators (see Table S2). Since the "DPL model" presented some instability, we further discuss outputs for this one specifically. Because we faced several issues when fitting the DPL model (including infants), we detail subsequently our rationale. Initially, "group" was to be considered as a random effect. However, doing so lead to violation of necessary assumptions (heteroscedasticity). This was due to the fact that group ID was partially nested into the group size (i.e. some groups could easily be identified based on their group size. For these groups, group size was indeed higher than for the other, and with lower variation along time). As such, we chose to consider group as a fixed effect. It solved heteroscedasticity issues, but induced weak correlation issue (max VIF $> 3$, see Table S2). We considered this less problematic than the first situation [Schielzeth et al., 2020].

Because infants of less than 1 yo are highly dependent on their mother for feeding, but also for moving, we considered group size when counting them or not. In this supplementary file, we provide a visual representation of differences between the two fitted DPL models (see Figure S6).

In the end, we observed no major deviation from necessary assumptions although we noticed statistically significant deviations (Figures, S8, S9). This might be in part induced by the skewed distribution of DPL. It is important, however, to note that the test for normality (Kolmogorov-Smirnov), as well as for heteroscedasticity of the *DHARMa* package are extremely sensitive (see CRAN vignettes of the package). In addition, these minor deviations could be imputed to the VIF troubles, and we discuss below how we further investigated the overall model stability.

We also investigated the sensibility of models to singular observations (i.e. withdrawal of one observation at a time) and to random levels (i.e. withdrawal of one level at a time, represented by the "month" variable) when accounting or not for infants of less than one year using the *influence.ME* package [Nieuwenhuis et al., 2012]. We checked the amplitude of the variations of estimates (Table S2).

#### 5.4 The relationships between group size and infants number

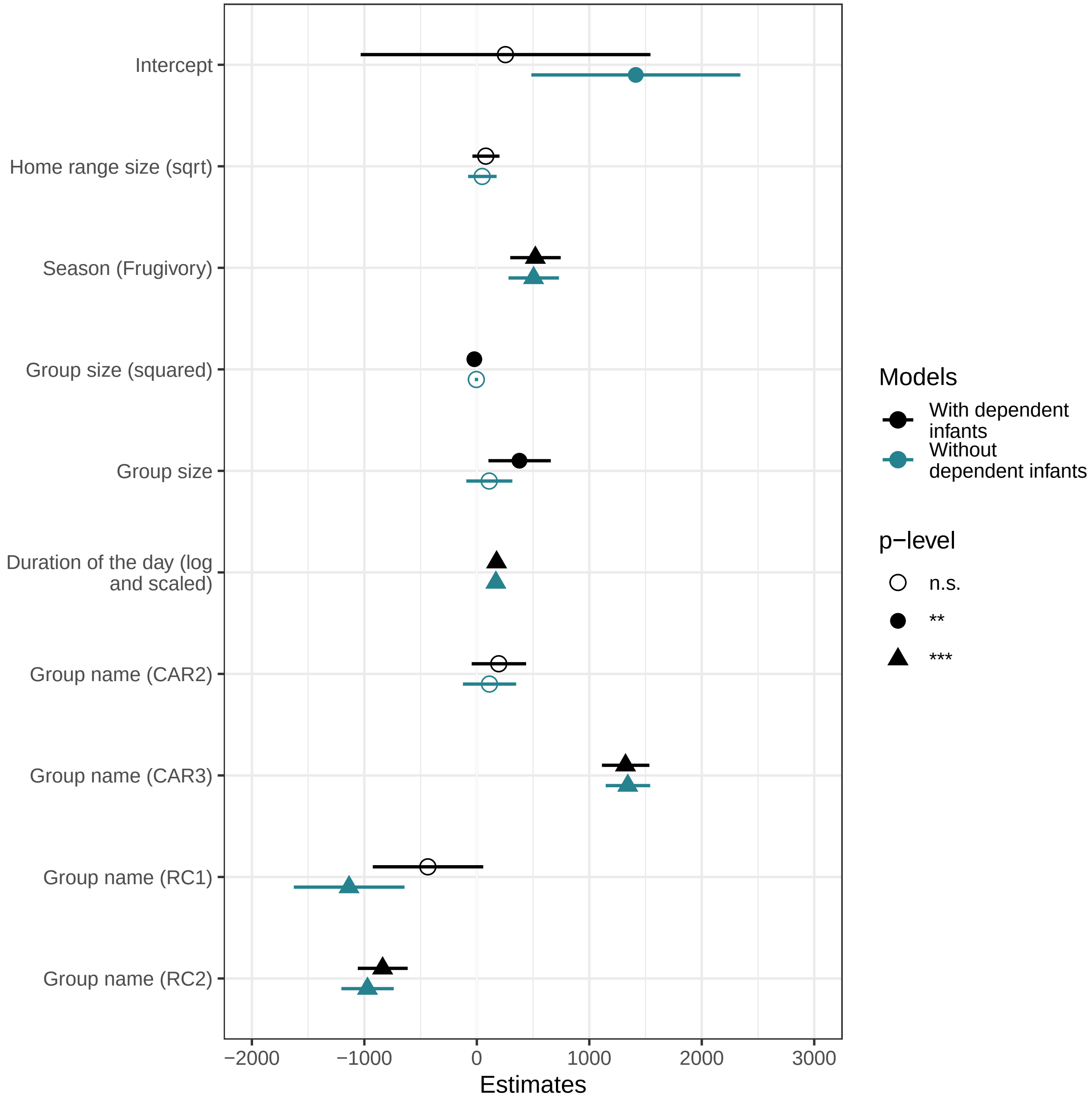


Figure S6: Forest plot of DPL model estimates when including or excluding infants

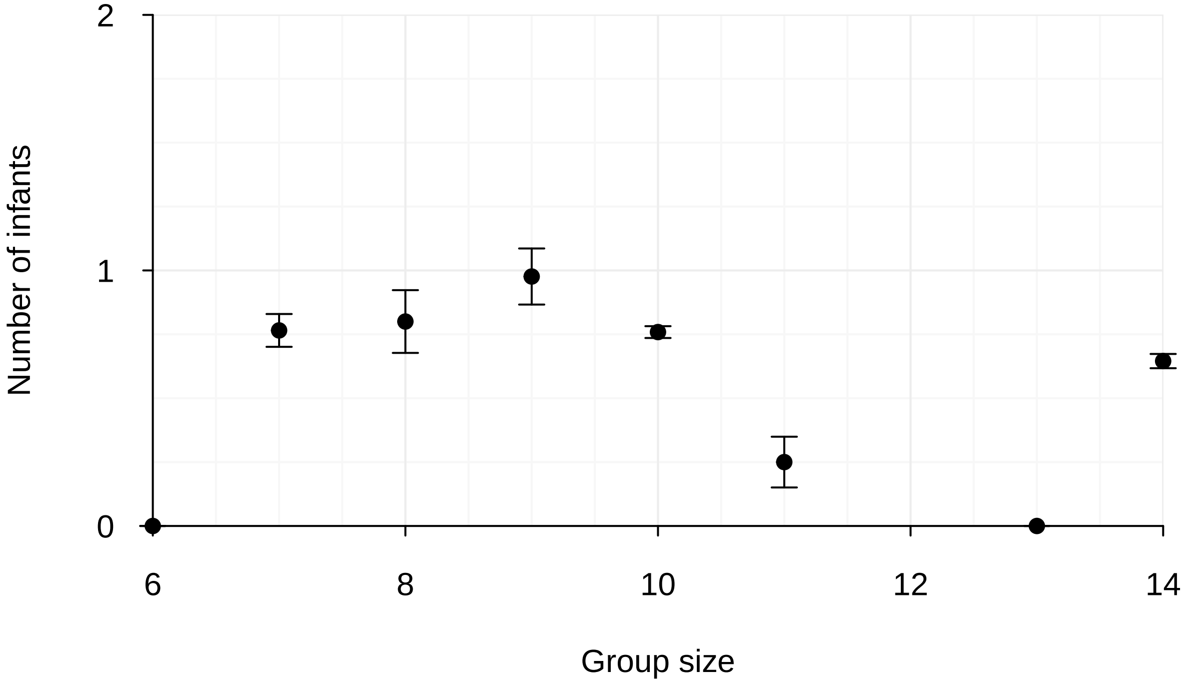


Figure S7: Number of infants as a function of group size | Means and standard errors are represented by points and vertical segments.

### 6 Model assumptions, stability and influence measures

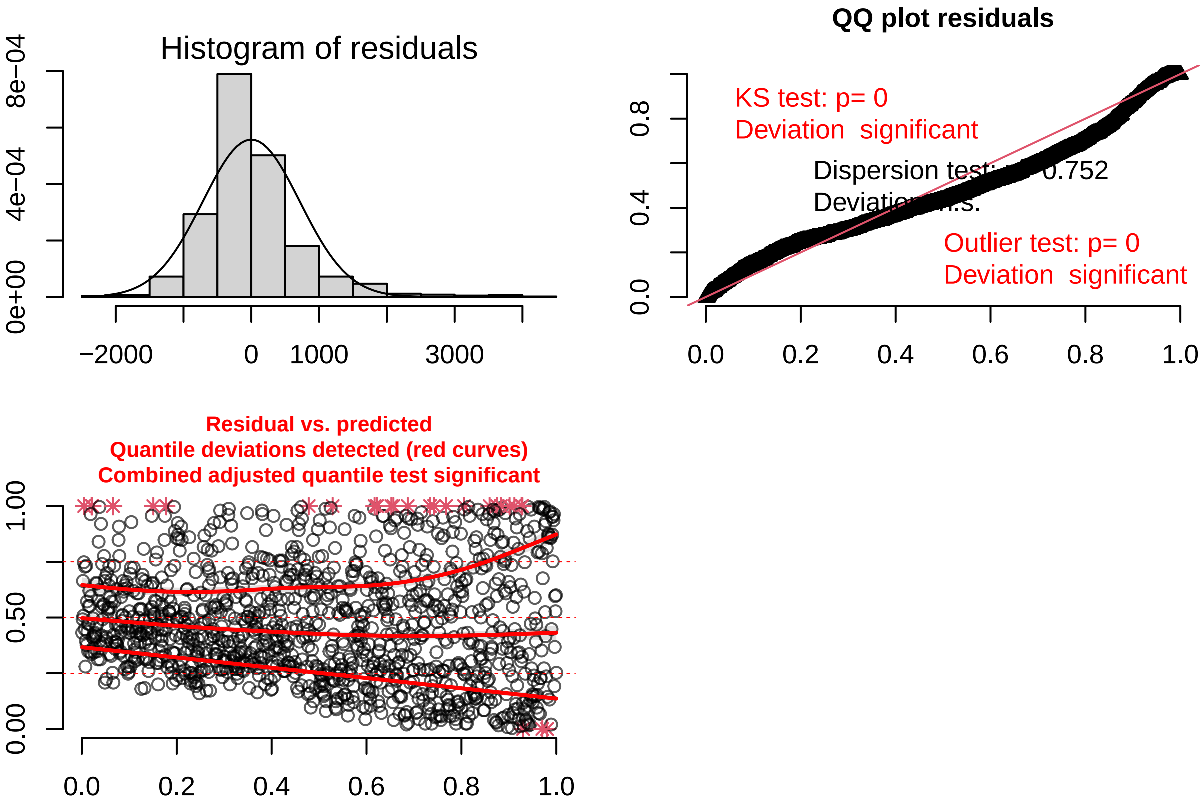


Figure S8: Model assumptions check: 'DPL' model with infants | The histogram of residuals is shown, as well as the Q-Q plot (adapted for the modelled distribution; with test for deviation of the distribution (Kolmogorov-Smirnov, KS, outliers and overdispersion) and the scatter plot of the fitted values vs. the residuals. Plots were based on the *DHARMa* package [Hartig, 2022].

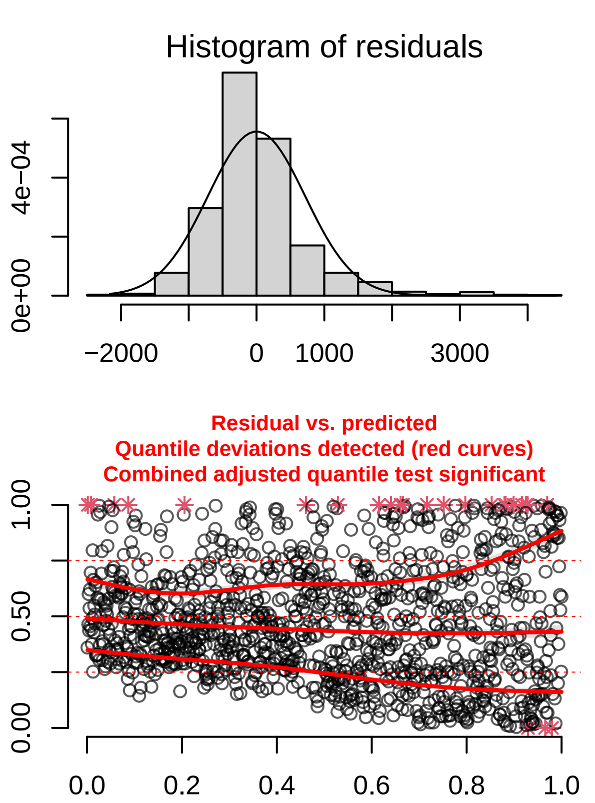


Figure S9: Model assumptions check: 'DPL' model without infants | The histogram of residuals is shown, as well as the Q-Q plot (adapted for the modelled distribution; with test for deviation of the distribution (Kolmogorov-Smirnov, KS, outliers and overdispersion) and the scatter plot of the fitted values vs. the residuals. Plots were based on the *DHARMa* package [Hartig, 2022].

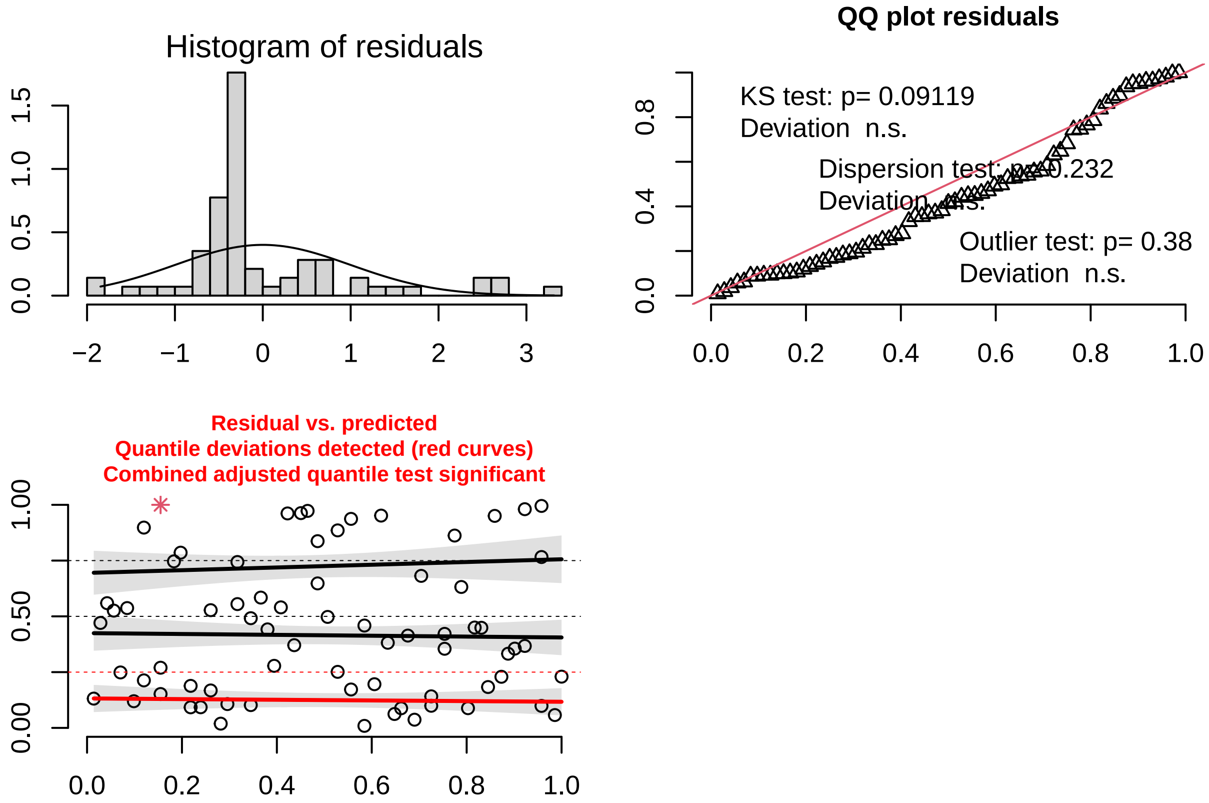


Figure S10: Model assumptions check: 'Visit timing' models (when AIC minimized) | The histogram of residuals is shown, as well as the Q-Q plot (adapted for the modelled distribution; with test for deviation of the distribution (Kolmogorov-Smirnov, KS, outliers and overdispersion) and the scatter plot of the fitted values vs. the residuals. Plots were based on the *DHARMa* package [Hartig, 2022].

### 7 Model stability and influence measures

Table S2: Dfbetas for all linear models | Obs. level shows the results when one singular observation was withdrawn at a time. Random level the shows results when one level of random effect ('month' for DPL models, 'group' for Visits model) was withdrawn at a time.

| Model: | | Obs. level | | Random level | |
|---|---|---|---|---|---|
| | Estimate | Min | Max | Min | Max |
| Model DPL: with infants Max VIF = 3.4 Cook's distance (obs. level) = 0.06 | | | | | |
| Intercept | 254.25 | 253.85 | 254.45 | 253.68 | 254.83 |
| Home range size (sqrt) | 79.55 | 79.4 | 79.77 | 78.28 | 80.02 |
| Season | 520.28 | 520.21 | 520.36 | 519.71 | 520.82 |
| Group size (squared) | -22.25 | -22.69 | -22.04 | -22.88 | -21.61 |
| Group size | 378.9 | 378.7 | 379.3 | 378.37 | 379.61 |
| Duration of the day (log and scaled) | 175.58 | 175.04 | 175.89 | 174.43 | 176.51 |
| Group name (CAR2) | 194.69 | 194.54 | 195.04 | 194.08 | 195.35 |
| Group name (CAR3) | 1321.89 | 1321.63 | 1322.18 | 1320.12 | 1322.6 |
| Group name (RC1) | -435.52 | -435.66 | -435.02 | -436.17 | -435.01 |
| Group name (RC2) | -837.73 | -837.95 | -837.4 | -838.36 | -837.16 |
| Model DPL: without infants Max VIF = 3.59 Cook's distance (obs. level) = 0.04 | | | | | |
| Intercept | 1412.96 | 1412.74 | 1413.23 | 1412.16 | 1414.21 |
| Home range size (sqrt) | 47.6 | 47.45 | 47.8 | 46.72 | 48.09 |
| Season | 504.68 | 504.61 | 504.75 | 504.13 | 505.25 |
| Group size (squared) | -4.6 | -4.77 | -4.35 | -5.19 | -3.71 |
| Group size | 109.75 | 109.51 | 109.92 | 108.71 | 110.66 |
| Duration of the day (log and scaled) | 169.43 | 168.88 | 169.72 | 168.34 | 170.35 |
| Group name (CAR2) | 111.85 | 111.68 | 112.17 | 110.22 | 112.74 |
| Group name (CAR3) | 1342.21 | 1341.94 | 1342.53 | 1340.39 | 1343.12 |
| Group name (RC1) | -1136.16 | -1136.33 | -1135.99 | -1137.28 | -1135.52 |
| Group name (RC2) | -972.35 | -972.6 | -972.15 | -974.15 | -971.64 |
| Model Visits Core Area Cook's distance (obs. level) = 0.22 | | | | | |
| Intercept | -3.69 | -3.78 | -3.36 | -4.75 | -2.7 |
| cos(month + best phase) | 0.43 | -0.21 | 1.03 | -0.81 | 0.95 |

### 8 Difference of visit duration within the neighbouring group heart area: check of t-test assumptions

We visually assessed deviation of residuals of normality and homoscedasticity based on the *DHARMa* package [Hartig, 2022] on linear models (response is visit duration, predictor is group ID), separately for each dyad. Models were fitted with the "lm" function of the *stats* package available together with the *base* package.

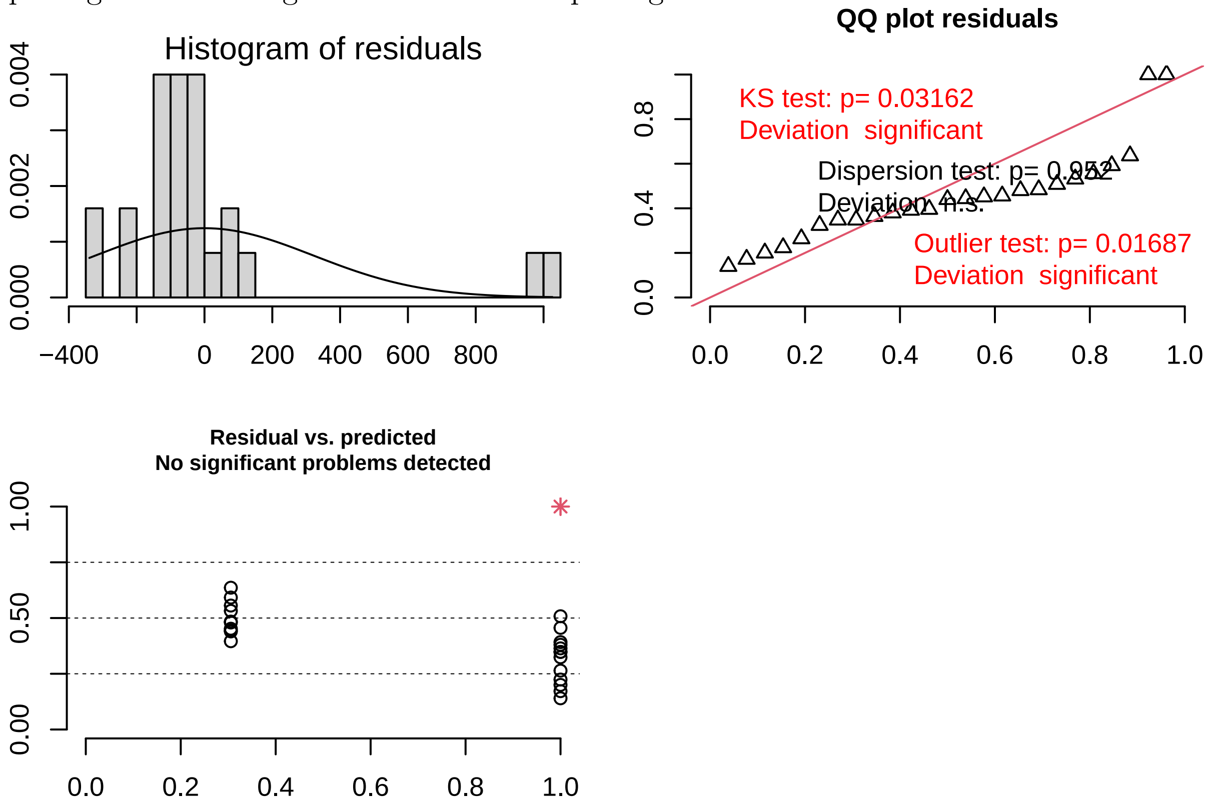

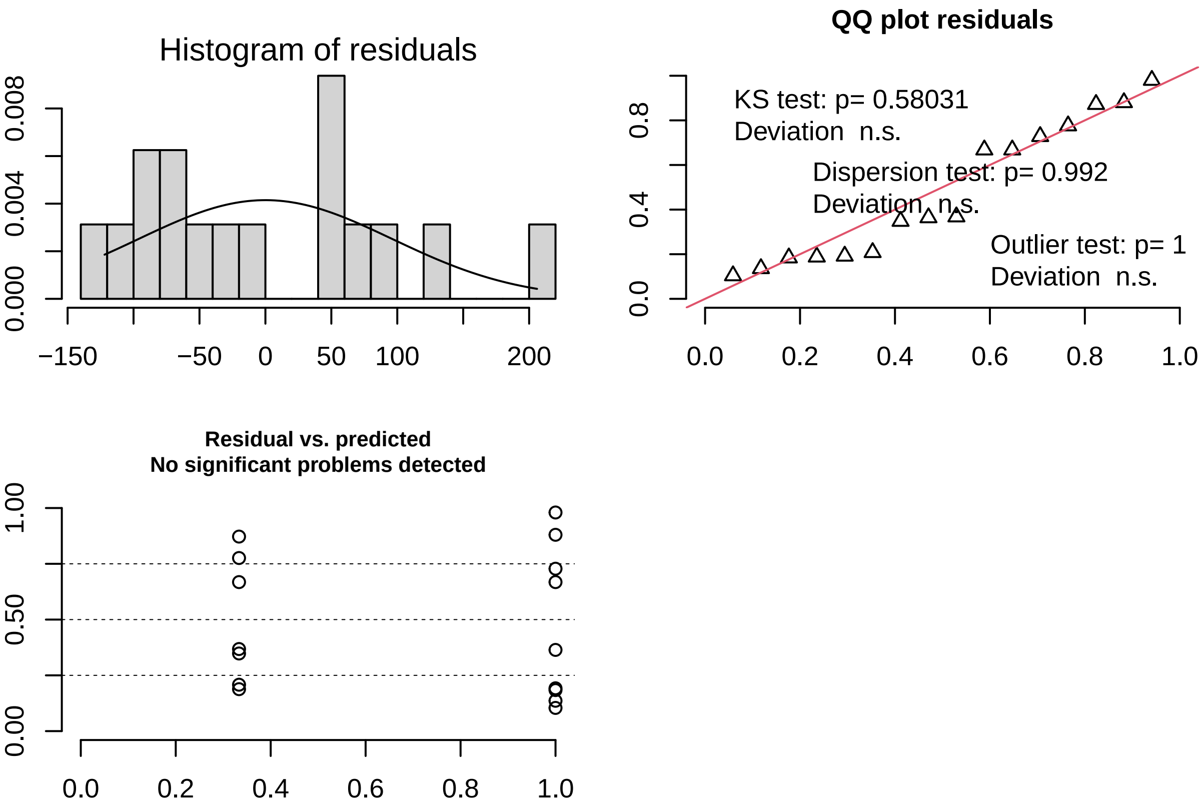
